# An ordered pattern of Ana2 phosphorylation by Plk4 is required for centriole assembly

**DOI:** 10.1101/240374

**Authors:** Tiffany A. McLamarrah, Daniel W. Buster, Brian J. Galletta, Cody J. Boese, John M. Ryniawec, Natalie A. Hollingsworth, Amy E. Byrnes, Christopher W. Brownlee, Kevin C. Slep, Nasser M. Rusan, Gregory C. Rogers

## Abstract

Polo-like kinase 4 (Plk4) initiates an early step in centriole assembly by phosphorylating Ana2/STIL, a structural component of the procentriole. Here, we show that Plk4 binding to the central coiled-coil (CC) of Ana2 is a conserved event, involving Polo-box 3 and a previously unidentified putative CC located adjacent to the kinase domain. Ana2 binding stimulates Plk4 kinase activity in vitro, and, in turn, is phosphorylated along its length. Previous studies showed that Plk4 phosphorylates the C-terminal STAN domain of Ana2/STIL, triggering binding and recruitment of the cartwheel protein Sas6 to the procentriole assembly site. However, the physiological relevance of N-terminal phosphorylation was unknown. We found that Plk4 first phosphorylates the extreme N–terminus of Ana2 which is critical for subsequent STAN domain modification. Phosphorylation of the central region then breaks the Plk4-Ana2 interaction. This phosphorylation pattern is important for centriole assembly and integrity because replacement of endogenous Ana2 with phospho-Ana2 mutants disrupts distinct steps in Ana2 function and inhibits centriole duplication.

## Introduction

Centriole duplication begins with the formation of a procentriole, which assembles orthogonally from the proximal end of a parent centriole (Fu et al., 2015). Procentriole assembly involves the hierarchical recruitment of a conserved set of proteins, SPD-2/DSpd-2/Cep192, Plk4/ZYG-1, SAS-5/Ana2/STIL, Sas6 and Sas4/CPAP, to a single assembly site on the mother centriole (Avidor-Reiss and Gopalakrishnan, 2013). In addition, the activity of the regulatory kinase, Polo-like kinase 4 (Plk4), is essential for centriole assembly, and Plk4 overexpression induces not only amplification from pre-existing centrioles but also de novo assembly (Bettencourt-Dias et al., 2005; Habedanck et al., 2005; Peel et al., 2007; Rodrigues-Martins et al., 2007; Kleylein-Sohn et al., 2007; Holland et al., 2010; Lopes et al., 2015). Characterizing Plk4 regulation of specific substrates is key to understanding centriole biogenesis.

*Drosophila* Anastral Spindle 2 (Ana2; STIL in humans) is an essential core centriole protein that follows Plk4 to the procentriole assembly site, and contains an N-terminal Sas4–binding site, a central coiled-coil (CC), and a C-terminal STAN domain (Fig. S1 A) (Goshima et al., 2007; Stevens et al., 2010a; Tang et al., 2011; Cottee et al., 2013; Hatzopoulos et al., 2013). Ana2 tetramerizes through the CC and binds Sas6, a rod-shaped protein, through the STAN domain (Stevens et al., 2010a; Shimanovskaya et al., 2013; Dzhindzhev et al., 2014; Slevin et al., 2014; Cottee et al., 2015). Ana2 tetramers may bind and facilitate Sas6 assembly into rings on the mother centriole’s surface, ultimately creating the stack of Sas6 rings that form the cartwheel of the nascent procentriole (Stevens et al., 2010b; Guichard et al., 2012; Dzhindzhev et al., 2014; Cottee et al., 2015; Moyer et al., 2015; Rogala et al., 2015).

Plk4 extensively phosphorylates Ana2/STIL, especially the STAN domain which promotes Sas6 binding (Dzhindzhev et al., 2014; Ohta et al., 2014; Kratz et al., 2015; Moyer et al., 2015), a critical event that is required for Sas6 recruitment to procentrioles. In addition, Plk4 phosphorylates several upstream residues in Ana2/STIL with unknown functional consequences (Dzhindzhev et al., 2014; Ohta et al., 2014; Kratz et al., 2015). Plk4 also binds the coiled-coil (CC) domain in STIL (Ohta et al., 2014; Moyer et al., 2015; Kratz et al., 2015), utilizing Plk4’s PB3 domain and an obscure second site located within its linker (L1) region (Arquint et al., 2015). Current models suggest that Ana2 is recruited to centrioles through its interaction with Plk4, as CC deletion mutants prevent Plk4 binding and centriole targeting (Ohta et al., 2014; Arquint et al., 2015; Cottee et al., 2015; Moyer et al., 2015). Our results here suggest that Ana2 binds Plk4 at two distinct sites, and that binding relieves Plk4 autoinhibition and activates the kinase. In turn, Plk4 phosphorylates clusters of Ana2 residues in an ordered pattern, and these modifications are critical for centriole assembly and integrity in cells. Furthermore, our findings suggest that Ana2 localization to centrioles can occur without direct binding to Plk4.

## Results

### Ana2 interacts with two distinct regions of Plk4

To test whether Plk4 and Ana2 binding occurs in *Drosophila*, we co-expressed transgenic V5- Ana2 and Plk4-GFP in S2 cells depleted of endogenous Ana2 (Fig. S1 B and C). Because Ana2 oligomerizes, we eliminated the influence of endogenous Ana2 on the binding assay by targeting its UTR with RNAi. Whereas full-length (FL) Ana2 co-immunoprecipitated (co-IP) with Plk4, Ana2 lacking the CC (ΔCC) did not (Fig. S1 D). Thus, the Plk4-Ana2 interaction is conserved in flies and requires the Ana2-CC.

Plk4 contains several functional domains (Klebba et al., 2015a): a N-terminal kinase domain followed by the Downstream Regulatory Element (DRE), and three Polo-boxes (PB1-3) interrupted by linkers L1 and L2 (Fig. 1 A). Previous efforts to map the STIL binding domain in human Plk4 have generated conflicting results and implicate multiple sites of interaction (Ohta et al., 2014; Arquint et al., 2015). We performed co-IP experiments using V5-Ana2 with either FL or truncated Plk4-GFP proteins to map Ana2 binding sites (Fig. 1 A). Ana2 associates with Plk4-FL and PB3 but not PB1-PB2 (Fig. 1 B), in agreement with Arquint et al. (2015). Ana2 also weakly co-IPs with Plk4 1-381, which lacks PBs. Yeast two-hybrid analysis confirmed these results: Ana2 interacts with 1-381 and PB3, but not PB1-PB2 (Fig. S1 E). Therefore, in contrast to a recent in vitro study indicating that Ana2 does not bind PB3 (Cottee et al., 2017), our results suggest that Ana2 associates with PB3 as well as an N-terminal region restricted to 1-381. The possibility that Plk4 kinase activity regulates its interaction with STIL has been investigated but remains unresolved (Ohta et al., 2014; Moyer et al., 2015). We examined this in S2 cells by co-expressing Ana2 with either wild-type (WT) or kinase-dead (KD) Plk4 truncation fragments 1-317 and 1-381 and then evaluating their interactions by co-IP. Consistently, Ana2 associated more with inactive Plk4-KD than WT (Fig. 1 C), suggesting that kinase activity suppresses this interaction in *Drosophila*. Furthermore, when combined with our finding that purified Plk4 1-317 binds GST-Ana2 using an in vitro pulldown assay (Fig. S1 F), these results also demonstrate that L1 is not necessary for Ana2 binding.

**Figure 1.**
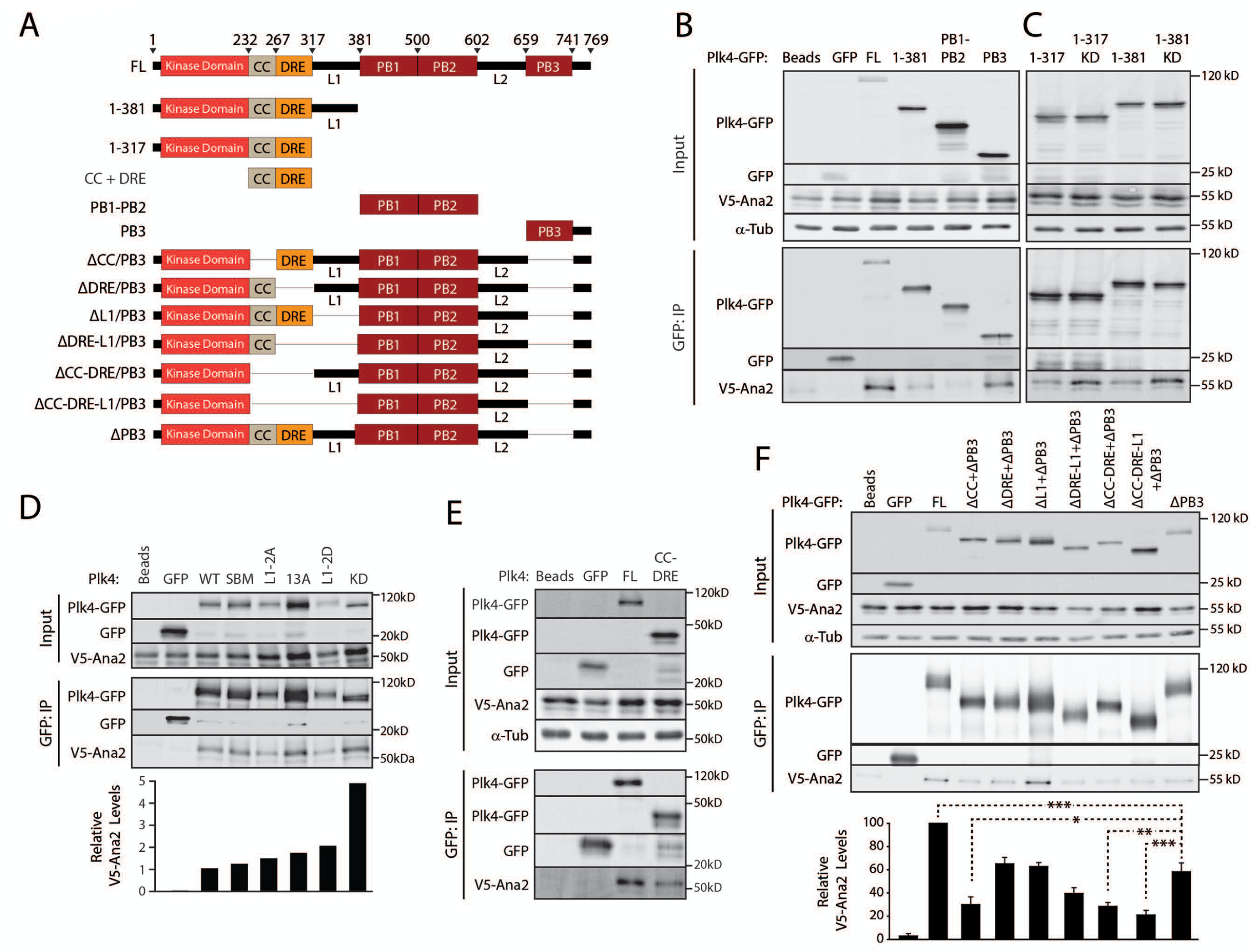
Ana2 binds both the putative CC-DRE and PB3 domains in Plk4. (A) Plk4 constructs used in immunoprecipitation (IP) experiments. CC, coiled-coil; DRE, Downstream Regulatory Element; PB, Polo Boxes; L1 and L2, linkers. (B) Ana2 associates with Plk4 N-terminus (1-381) and C-terminus (PB3). S2 cells were cotransfected with Plk4-GFP [full-length (FL) or deletion constructs] and V5-Ana2. Anti-GFP IPs were performed from lysates of the transfected cells and probed for GFP, V5, and α-tubulin. (C) L1 is not necessary to associate with Ana2, and Ana2 interacts preferentially with kinase-dead (KD) Plk4. IPs performed as in B. (D) Ana2-Plk4 binding is greatest when the kinase is inactive (KD). Plk4 constructs evaluated are: WT, wild-type; SBM, non-degradable Slimb-binding mutant; L1-2A and 2PM, contains two Plk4-targeted serines of L1 mutated to alanine (A) or phosphomimetic (PM) residues; 13A, contains 13 Plk4-targeted DRE residues mutated to alanine. Each relative Ana2 level measured using the integrated intensity of a bound Ana2 band relative to its corresponding immunoprecipitated Plk4-GFP band. Graph shows relative-Ana2 levels normalized to the WT result. (E) The Plk4 CC-DRE domain is sufficient to associate with Ana2. IPs performed as in B. (F) Ana2 associates with the putative CC and PB3 regions of Plk4. Graph depicts relative intensity of V5-Ana2 normalized to Plk4-GFP in the IP. Asterisks mark significant differences between treatments. Error bars, SEM; n = 3 independent experiments.

One possible explanation for an enhanced interaction of Ana2 with Plk4-KD is that Plk4 autophosphorylation prevents Ana2 binding. To test this, we mutated residues in the DRE and L1 regions known to be autophosphorylated (Klebba et al., 2013; Cunha-Ferreira et al., 2013; Klebba et al., 2015a), and examined Ana2 association by IP. Ana2 co-IPed with both WT-Plk4 and a non-degradable Slimb-binding mutant (SBM) (Fig. 1 D). Similar levels of Ana2 associated with Plk4 when 13 serines in the DRE are mutated to non-phosphorylatable alanines (13A), as well as when 2 serines of L1 are mutated to either non-phosphorylatable alanine or phosphomimetic aspartic acid (L1-2A/2D). Notably, Ana2 interacted with full-length inactive Plk4-KD at approximately 5-fold higher levels than WT, indicating that Plk4 kinase activity disrupts Ana2 association, and that this effect is not due to autophosphorylation of the DRE or L1.

Using protein structure prediction software, we identified a conserved, previously uncharacterized coiled-coil (CC) within Plk4 1-317, immediately adjacent to the kinase domain (Fig. S1 G, H). To test the role of the putative Plk4-CC in Ana2 binding, we performed co-IPs from S2 cells expressing Plk4 constructs lacking PB3 and one or more N-terminal modules (Fig. 1 F). Ana2 association was decreased ~2-fold by deletion of PB3 and was significantly reduced a further ~2-fold by deletion of both the Plk4 putative CC and PB3 regions. Combined with the observation that Ana2 associates with a Plk4 fragment consisting of only CC-DRE (Fig. 1 E), this result pinpoints the putative CC of Plk4 as the other Ana2-binding domain besides PB3, although we cannot rule out additional Ana2-binding domains.

### Ana2 stimulates Plk4 kinase activity in vitro

STIL overexpression stimulates Plk4 kinase activity in cultured human cells (Ohta et al., 2014; Moyer et al., 2015), and it has been proposed that STIL binding stimulates Plk4 activity by relieving L1-mediated autoinhibition (Arquint et al., 2015). In this model, STIL binds both PB3 and L1, and repositions L1 so that it no longer prevents Plk4 from autophosphorylating its activation loop. This model integrates well with our previous finding that L1 reduces Plk4 kinase activity (Klebba et al., 2015a). To test this model further, we first determined if Ana2 stimulates Plk4 activity by mixing purified full-length His-MBP-Plk4 with either GST or GST-Ana2 and then measuring Plk4 autophosphorylation. Without Ana2, Plk4 displayed a noticeable lag before reaching its maximal rate of autophosphorylation, giving rise to a sigmoidal plot that is expected for a *trans*-activating kinase (Fig. 2 A), as was also shown by Lopes et al. (2015). However, in the presence of Ana2, the kinetics of Plk4 autophosphorylation shifted to a hyperbolic curve demonstrating rapid Plk4 activation (Fig. 2 A). (Purified GST-Ana2 alone did not have detectable kinase activity [Fig. S2 A]).

**Figure 2.**
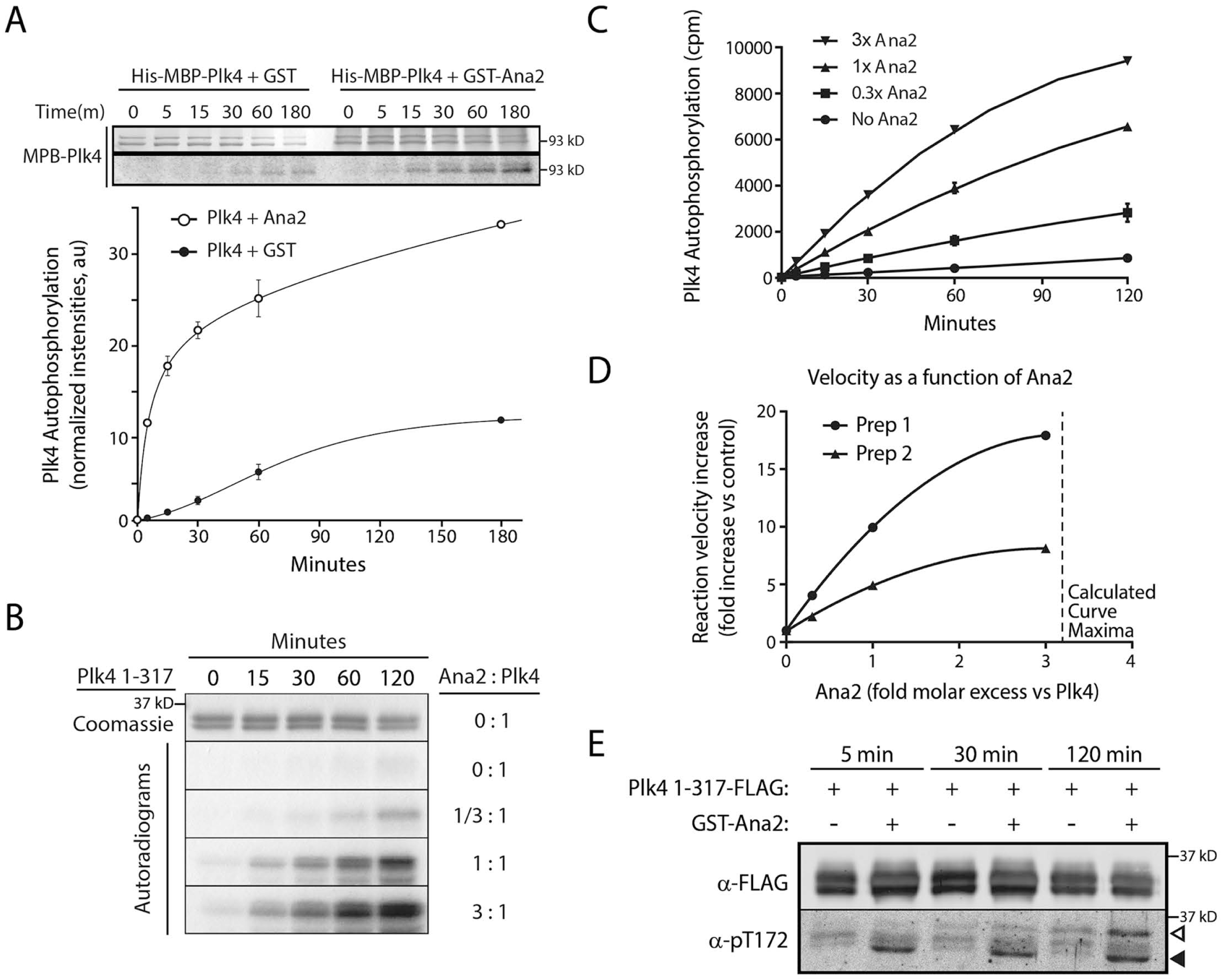
Ana2 stimulates Plk4 autophosphorylation in vitro. (A) Ana2 stimulates full-length Plk4 activity in vitro. Purified full-length His-MBP-Plk4 was mixed with equimolar amounts (2.5 μM) of either GST or GST-Ana2 and incubated with γ^32^P-ATP. Reactions were sampled at intervals (0–180 min). Top, Coomassie-stained gel; bottom, phosphorimage. Graph shows relative intensities of Plk4 autophosphorylation (normalized to Coomassie-stained bands) over time. In addition to the difference in initial rates, the calculated horizontal asymptotes (limits of autophosphorylation) of the two plots are different. Possibly, the presence of Ana2 allows Plk4 to autophosphorylate sites that are normally not accessible. *n* = 2 experiments. Error bars, SEM. (B) Purified Plk4 1-317-His was incubated with γ^32^P-ATP and different concentrations of purified full-length GST-Ana2. Reactions were sampled at intervals (0–120 min). Top panel, Coomassie-stained Plk4 1-317; 4 bottom panels, autoradiograms. The molar ratios of Ana2 to Plk4 in each assay are indicated on the right. (C) Ana2 significantly increases the rate of Plk4 autophosphorylation. Bands of Plk4 1-317 from samples resolved by SDS-PAGE were cut from gels, and their ^32^P-radiolabel measured by scintillation counting. The initial rates for all 4 plots are significantly different (P<0.001). (D) The initial rates of autophosphorylation for two different Plk4 1-317 preparations were examined as in B, and the fold increase in rate as a function of Ana2 plotted. The maximum for each best-fit plot is reached when the Ana2 molar concentration exceeds Plk4 by 3–4x. (E) GST-Ana2 increases the autophosphorylation of activation loop residue T172 in Plk4 1-317. Purified Plk4 1-317-His was incubated with 3x molar GST-Ana2 and ATP, and the mixture sampled at the indicated time points. Western blots were probed with anti-FLAG (to detect all Plk4) and anti-phospho-T172 antibodies. Arrowheads mark 2 prominent pT172-positive species of Plk4 that appear in the presence of Ana2.

If Ana2 stimulates Plk4 activity by relieving L1-mediated autoinhibition, then Ana2 should fail to stimulate Plk4 lacking L1. To test this, purified minimal Plk4 protein (amino acids 1-317) -- containing the kinase domain-CC-DRE but lacking L1 -- was assayed in vitro in the presence of increasing concentrations of GST-Ana2. Plk4 1-317 autophosphorylation was low in the absence of Ana2, but surprisingly, increased with addition of Ana2 in a dose-dependent manner (Fig. 2 B, C). A plot of the initial reaction velocities as a function of Ana2 concentration indicates that autophosphorylation rate is maximal when Ana2 is in 3–4x molar excess to Plk4(Fig. 2 D). We also examined the effect of Ana2 on a longer Plk4 construct (amino acids 1-602) containing L1 and PB1-PB2. As expected, Plk4 1-602 autoinhibits, displaying a low level of kinase activity despite being able to dimerize (Fig. S2 B), as previously described (Klebba et al., 2015a). Notably, Ana2 failed to stimulate the kinase activity of the autoinhibited 1-602 construct as it did the 1-317 construct (Fig. S2 B, C), suggesting that the PB3 domain is required for Ana2 to accelerate Plk4 autophosphorylation when L1 is present.

Plk4 activity is greatly increased by *trans*-autophosphorylation of its activation loop, particularly the conserved threonine, T172 (Swallow et al., 2005; Klebba et al., 2015a; Lopes et al., 2015). If Ana2 stimulates Plk4 kinase activity, then T172 phosphorylation should also increase. To test this, we generated antibodies specific to phosphorylated T172 (Fig. S2 D) and found that phosphorylated T172 (pT172) in Plk4 1-317 either increased or decreased depending on the presence or absence, respectively, of Ana2 (Fig. 2 E). Interestingly, anti-pT172 detected two prominent bands of activated Plk4: an upper, ostensibly hyper-phosphorylated band (open arrowhead) and a lower, less phosphorylated band (filled arrowhead). Since prominent anti-pT172 staining first appears in the lower band, T172 may be one of the first autophosphorylated residues. Taken together, our results suggest that Ana2 stimulates Plk4 activity utilizing a two-step mechanism. First, Ana2 binds both the putative CC and PB3 in Plk4 and, by manipulating L1, relieves Plk4 autoinhibition. In the second step, Ana2 then stimulates Plk4 kinase activity by a mechanism independent of L1; specifically, we propose that tetrameric Ana2 binds multiple Plk4s (likely through the putative CC) in a proximity and orientation that facilitate *trans*-autophosphoryl ati on.

### Plk4 phosphorylation of Ana2 residues upstream of the STAN domain is important for centriole duplication

Previous studies show that Ana2/STIL is phosphorylated by Plk4 and that modification of the STAN domain promotes Sas6 binding (Dzhindzhev et al., 2014; Ohta et al., 2014; Kratz et al., 2015; Moyer et al., 2015). We used tandem mass spectrometry (MS/MS) to identify phosphorylated residues of Ana2 incubated with Plk4 in vitro. Plk4 phosphorylated Ana2 on eleven residues (Fig. 3 A, Table S1A): four residues within the STAN domain and seven upstream residues that displayed varying conservation across phyla (Fig. S3 A). Plk4 is not a highly promiscuous kinase; e.g., it does not phosphorylate Sas6 (Fig. S3 B) (Dzhindzhev et al., 2014). Notably, seven of the residues we mapped were previously shown to be either phosphorylated in cells co-expressing Plk4 or identified from samples mixed with Plk4 in vitro (Dzhindzhev et al., 2014). We also examined in vivo phosphorylation by isolating transgenic GFP-Ana2 from asynchronous S2 cells and performing MS/MS (Table S1 B). Although our peptide coverage lacked amino acids 1-65, we found that Ana2 is phosphorylated on numerous residues, including six residues also identified in vitro (Fig. S3 C).

**Figure 3.**
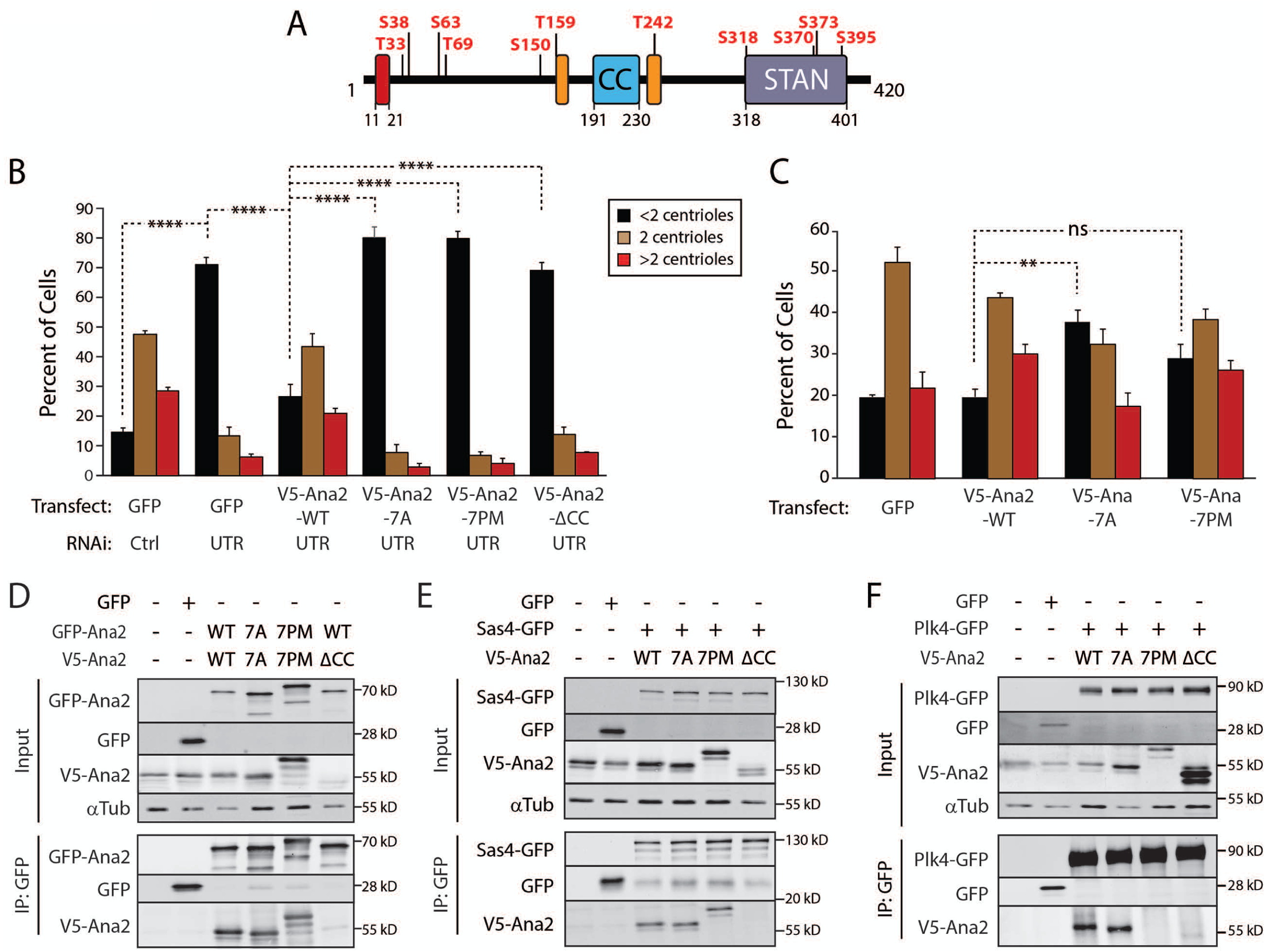
Ana2 is a Plk4 substrate. Expression of phospho-Ana2 mutants inhibit centriole duplication. (A) Linear map of Ana2 depicting the Plk4 phosphorylation sites. Phospho-sites were identified by tandem MS of purified GST-Ana2 phosphorylated in vitro by purified Plk4 1-317-His_6_. Red box, Sas4-binding site (Cottee et al., 2013); orange boxes, LC8 binding-sites (Slevin et al., 2014). (B) N-terminal phosphorylation mutations in Ana2 disrupt centriole duplication. On days 0, 4, and 8, S2 cells were control-dsRNA treated or depleted of endogenous Ana2 by RNAi (UTR). On days 4 and 8, cells were transfected with the indicated V5-Ana2 or GFP construct and then induced to express with 0.1 mM CuSO4. Cells were immunostained for PLP and Asterless to mark centrioles, and the number of centrioles per cell was counted. n = 100 cells in each of three experiments. Asterisks indicate significant differences. Error bars, SEM. (C) Expression of non-phosphorylatable Ana2-7A acts as a dominant/negative and inhibits centriole amplification. V5-Ana2 or GFP expression constructs were transfected on days 0 and 4 and induced with 0.5 mM CuSO4. Cells were fixed and immunostained on day 7. The average percentages of cells containing the indicated number of centrioles are shown (n = 100 cells in each of three experiments). Error bars, SEM. Asterisks indicate significant difference; ns, not significantly different. (D-F) The phosphorylation state of the Ana2 N-terminus does not affect Ana2 binding to itself (D) or Sas4 (E). However, phosphomimetic mutations in Ana2 (7PM) inhibit Plk4 binding, whereas Plk4 binding is not affected with non-phosphorylatable Ana2 (7A) (F). In contrast, the CC of Ana2 is required for Ana2 self-association (D) and binding to Sas4 and Plk4 (E, F). S2 cells were depleted of endogenous Ana2 by RNAi for 7 days. On day 5, cells were co-transfected with the indicated constructs and the next day induced to express for 24 hours. Anti-GFP IPs were then prepared from lysates, and Western blots of the inputs and IPs probed for GFP, V5, and α-tubulin.

Since the physiological consequence of Ana2/STIL phosphorylation outside the STAN domain is unknown, we focused on the seven upstream phospho-residues (T33/S38/S63/T69/ S150/T159/T242) that we mapped in vitro and examined their effects on centriole duplication. First, we generated non-phosphorylatable and phosphomimetic Ana2 by mutating all seven residues to alanines (7A) or aspartate/glutamate (7PM) and transfected these into S2 cells depleted of endogenous Ana2. Centriole numbers were counted after immunostaining for centriole-markers PLP and Asterless (Asl) (Mennella et al., 2012; Fu and Glover, 2012) (Fig. 3 B). As expected, Ana2 depletion significantly decreased the percentage of cells with a normal number of centrioles (Fig. 3 B). In cells lacking endogenous Ana2, centriole numbers were rescued by expression of Ana2-WT, but, surprisingly, not by Ana2-7A, Ana2-7PM, or Ana2 lacking the CC domain (ΔCC) which is required for both centriole assembly and localization (Dzhindzhev et al., 2014; Ohta et al., 2014; Arquint et al., 2015; Cottee et al., 2015; Moyer et al., 2015).

The failure of both Ana2-7A and 7PM to rescue centriole duplication could be explained if the substitutions cause protein misfolding and therefore prevent functionality, or if the two Ana2 phospho-mutants do fold properly but fail to support centriole duplication for unknown reasons. To evaluate these possibilities, we first purified recombinant Ana2-7A and 7PM and examined the structural stability of the mutant proteins versus the WT protein using circular dichroism (CD) (Fig. S3 D, E). No significant difference was found between the CD spectra of wild-type or mutant proteins, suggesting that the 7A/7PM substitutions were not altering the secondary structure of Ana2. Next, we expressed Ana2 constructs in untreated cells, where they can hetero-oligomerize with endogenous Ana2, and measured centriole numbers. Whereas Ana2-WT and Ana2-7PM expression had no effect on centriole number, Ana2-7A suppressed centriole duplication (i.e., it significantly increased the percentage of cells with less than 2 centrioles) (Fig. 3 C). Thus, unlike Ana2-7PM, Ana2-7A expression displays a dominant/negative effect on centriole duplication, suggesting that the Ana2-7A and 7PM mutants influence centriole assembly differently.

### Ana2 N-terminal phosphorylation by Plk4 blocks their association

We next asked whether the phosphorylation state of the Ana2 N-terminus alters Ana2’s interaction with known binding partners, including itself. Ana2 forms a tetramer through its CC, and point mutations that block oligomerization disrupt Ana2 function in vivo (Slevin et al., 2014; Cottee et al., 2015). We examined whether phospho-Ana2 mutants influence self-association using S2 cells depleted of endogenous Ana2. Ana2-ΔCC did not co-IP with Ana2-WT (Fig. 3 D), demonstrating that the CC is required for self-association, as it is for STIL (Arquint et al., 2015). However, both Ana2-7A and 7PM were able to co-IP with themselves, suggesting that the N-terminal phosphorylation state of Ana2 does not influence its oligomerization.

Since Sas4 binds Ana2 (Cottee et al., 2013; Hatzopoulos et al., 2013), another possibility is that Ana2 phosphorylation could affect this interaction. Sas4 associated with Ana2 regardless of its phosphorylation state (Fig. 3 E), consistent with previous in vitro studies (Dzhindzhev et al., 2014). However, Ana2-ΔCC association with Sas4 was markedly diminished, a surprising result since the Sas4-binding site in Ana2 is well upstream of the Ana2-CC (Fig. S1 A). Thus, Ana2 oligomerization may be a prerequisite for Sas4 binding.

Last, we examined the interaction between Ana2 and Plk4 (Fig. 3 F). Both Ana2-WT and Ana2-7A co-IPed with Plk4. Strikingly, however, Plk4 failed to IP with Ana2-7PM, similar to Ana2-ΔCC. In summary, our results reveal a complex relationship between Ana2 and Plk4: Ana2 binds Plk4 and increases Plk4 kinase activity, but when Ana2 is itself phosphorylated by Plk4, binding with Plk4 is inhibited. Thus far, our findings cannot explain how Ana2-7A acts as a dominant/negative of centriole duplication, but the inability of Ana2-7PM to bind Plk4 may explain why cells expressing this mutant cannot duplicate centrioles.

### The recruitment of Ana2-7A to procentrioles is disrupted but Ana2-7PM is not

Ana2 phospho-mutants can self-associate, bind Sas4 and, in the case of Ana2-7A, bind Plk4, suggesting that N-terminal substitutions do not cause protein misfolding in cells but instead interfere with specific functions of Ana2 that compromise centriole duplication. To further test this hypothesis, we used super-resolution (SR-SIM) microscopy to examine the centriolar localization of GFP-Ana2 (WT or mutant) in cells depleted of endogenous Ana2. In interphase cells, Ana2 staining typically appears as two spots that co-localize with a PLP-labeled mother centriole ring: a bright spot positioned near the center of the PLP ring and a second dimmer spot near the PLP periphery (Fig. 4 A-C, *top panels*) (Dzhindzhev et al., 2014). The dim Ana2 spot is recruited after telophase disengagement of a centriole pair and marks the future site of procentriole assembly by recruiting Sas6 (Dzhindzhev et al., 2014). In cells whose endogenous Ana2 was replaced with transgenic Ana2-WT, 75% of interphase centrioles (n=44) displayed the expected 2-spot Ana2 pattern; only 14% contained a single Ana2 spot within the mother (Fig. 4 A). Previous studies have shown that the CC in Ana2/STIL is necessary for centriole localization, perhaps by directly binding Plk4 (Ohta et al, 2014; Arquint et al., 2015; Cottee et al., 2015; Moyer et al., 2015). Since Ana2-7PM fails to associate with Plk4 even though it has an intact CC, we predicted that Ana2-7PM would fail to localize to the procentriole site, and instead appear only within the mother. Surprisingly, Ana2-7PM localized in a normal two puncta centriole pattern (74%, n=23) (Fig. 4 B). This suggests that Ana2 targets centrioles by a means other than direct Plk4 binding. In contrast, Ana2-7A localized as a single spot within the majority of mother centrioles (56%, n=39), and the expected two spot Ana2 pattern was observed in only 21% of mothers (Fig. 4 C). Thus, in most cells, Ana2-7A recruitment to the procentriole site fails, providing a possible explanation for why Ana2-7A expression prevents centriole duplication.

**Figure 4.**
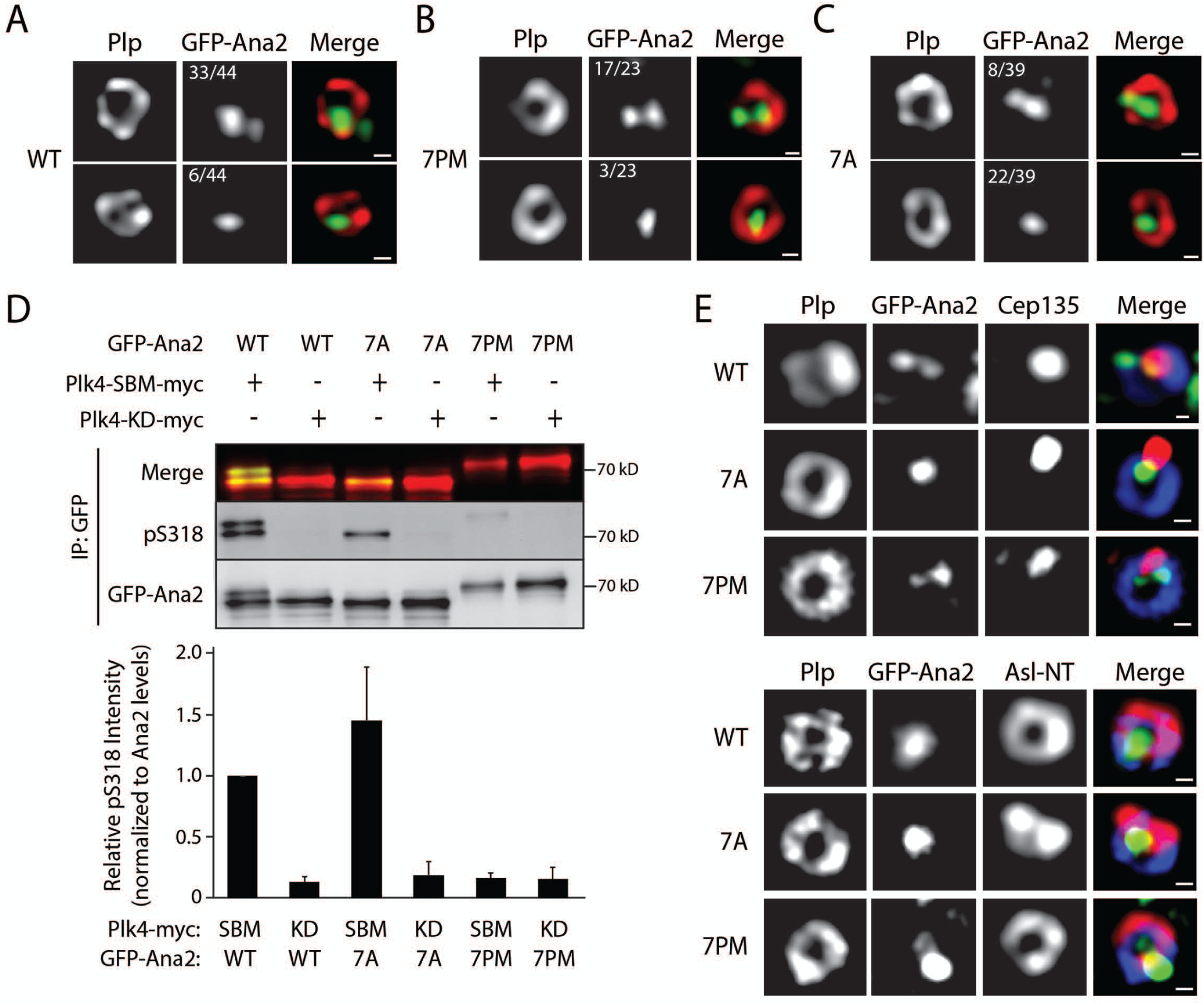
The phosphorylation state of the Ana2 N-terminus affects Ana2 recruitment to procentrioles and STAN domain phosphorylation. (A-C) Ana2-7A recruitment to procentrioles is inhibited but Ana2-7PM is not. Interphase centrioles in transgenic Ana2-expressing cells were imaged using super-resolution microscopy. S2 cell-lines stably transfected with inducible GFP-Ana2 WT, 7PM or 7A were depleted of endogenous Ana2 by RNAi (UTR) for 12 days and, during this period, were induced to express the indicated GFP-Ana2 (green) construct. Cells were immunostained with anti-PLP (red) to mark the surface of mature centrioles. DNA was visualized with Hoechst (not shown). The number of centrioles that show either two (upper panel) or one (lower panel) Ana2 spots is indicated. Centrioles containing either no detectable Ana2 or >2 spots comprised 11% (WT), 13% (7PM) and 23% (7A) of the remaining centrioles. Scale bars, 200 nm. (D) Phosphorylation of S318 in the STAN domain is strongly reduced by Ana2-7PM expression. Cells were depleted of endogenous Ana2 by RNAi for 7 days. On day 5, cells were cotransfected with the indicated GFP-Ana2 and Plk4-myc constructs and the next day induced to express for 24 hours. Anti-GFP IPs were then prepared from lysates, and blots of the IPs probed for GFP and pS318. A merge of GFP-Ana2 and pS318 blots is shown (upper panel). Graph shows relative pS318 intensities in the IPs. For each treatment, intensities of pS318 were measured, related to the corresponding GFP-Ana2 intensities, and then normalized to control (lane 1). n = 3 experiments. Error bars, SEM. (E) Asterless (Asl) distribution in centrioles containing Ana2-7A is abnormal. Interphase centrioles in Ana2-expressing cells imaged using super-resolution microscopy. Stable S2 cell lines were manipulated as in A. Cells were immunostained with Cep135 or Asl-NT (red), and PLP (blue). Hoechst-stained DNA not shown. Scale bars, 200 nm.

### STAN domain phosphorylation by Plk4 is reduced in Ana2-7PM

Although Ana2-7PM localizes normally to interphase centrioles, duplication is inhibited when Ana2-7PM replaces endogenous Ana2 in cells. Possibly, 7PM’s inability to bind Plk4 reduces phosphorylation of the STAN domain and, consequently, procentrioles cannot assemble. To test this, we generated a phospho-specific antibody against S318 (Fig. S3 F), a conserved STAN residue phosphorylated by Plk4 and required for Sas6 binding (Ohta et al, 2014; Dzhindzhev et al., 2014). In cells depleted of endogenous Ana2, GFP-Ana2 was co-expressed with either kinase-dead (KD) Plk4 or a highly active non-degradable Plk4-SBM. Western blots of GFP-Ana2 purified by IP from cell lysates were probed with the anti-phospho-S318 (pS318) antibody. This antibody recognized Ana2-WT that was co-expressed with active Plk4-SBM-myc but not with inactive Plk4-KD-myc (Fig. 4 D, lanes 1,2). Likewise, pS318 was detected in Ana2-7A co-expressed with active (but not inactive) Plk4 (Fig. 4 D, lanes 3, 4). In contrast, pS318 was almost entirely absent in Ana2-7PM, even when co-expressed with active Plk4 (Fig. 4 D, lanes 5, 6). Therefore, the presence of phospho-mimicking residues within the N-terminus of Ana2 prevents phosphorylation of the STAN domain (at least of S318) by Plk4 in cells. The reduction in STAN phosphorylation may account for the inability of Ana2-7PM to support centriole duplication. Taken together, our findings suggest that Ana2 binds Plk4 after being recruited to centrioles and this interaction is required for STAN phosphorylation. Interestingly, the pS318 antibody reacts with a tight doublet of Ana2 WT but only a single, fast-migrating band of Ana2-7A, which suggests that the seven substituted residues in the mutant are responsible for the electrophoretic shift of Ana2 WT’s second slower-migrating band. Unfortunately, the pS318 antibody recognizes additional polypeptides in blots of whole cell lysates and, therefore, was not useful for immunostaining studies.

### Centrioles containing Ana2-7A display morphological abnormalities

Thus far, our results do not explain the disruption of Ana2-7A localization to the procentriole site since Ana2-7A retains several WT features, such as oligomerization, binding of Sas4 and Plk4, and phosphorylation of its STAN domain (specifically, S318). An untested possibility is that proper localization of other procentriole-targeted proteins is altered when Ana2-7A is expressed. To examine this, we raised antibodies against Cep135 and the Asl N-terminal domain (Fig. S3 G) to evaluate the localization of these centriole proteins in cells. In centriole cross-sections, Asl-NT is positioned adjacent to PLP on the centriole surface, whereas Cep135 localizes as a single spot near the centriole center (Mennella et al., 2012; Fu et al., 2016). After depleting endogenous Ana2 and expressing the GFP-Ana2 constructs, we examined interphase centrioles using SR-SIM and observed normal localization patterns for both Cep135 and Asl in Ana2-WT and 7PM expressing cells (Fig. 4 E). Curiously, although Cep135 formed a normal single spot within centrioles in Ana2-7A cells, Asl staining was distorted and collapsed within the centriole as adjoining spots instead of assuming a normal ring shape (Fig. 4 E). Thus, Ala substitution of the N-terminal phospho-sites of Ana2 causes the redistribution of at least two duplication-essential proteins, Ana2 and Asl, within centrioles.

Next, we imaged PLP to examine centriole architecture in Ana2-7A expressing cells. In this case, endogenous Ana2 was not depleted in order to preserve more centrioles for observation. Using SR-SIM, we imaged PLP-labeled centrioles in GFP-Ana2 WT and 7A expressing interphase cells. We restricted our analysis to centrioles that were oriented orthogonally to the image plane (i.e., had a clear cross-sectional aspect) in order to more accurately measure the diameter (or major axis) of the PLP ring. In GFP-Ana2-WT expressing cells, 82% of centrioles (36 of 44) displayed a normal PLP ring with quasi-nine-fold symmetry and an average diameter of 486 nm (Fig. 5 A, C). The average diameter of the PLP ring in cells expressing GFP-Ana2-7PM was similar (515 nm) (Fig. 5 C). Although normal PLP arrangements were observed in GFP-Ana2-7A expressing cells (45%), the majority displayed an unusual spot or stripe of PLP in the centriole center (55%, 23 of 42) (Fig. 5 B). In addition, the average length of the major axis of PLP immunostaining (571 nm) was significantly greater than GFP-Ana2-WT-containing centrioles (Fig. 5 C). Despite these differences, centrioles containing GFP-Ana2-7A were functional in that they recruited γ-tubulin during mitosis (Fig. S3 H).

**Figure 5.**
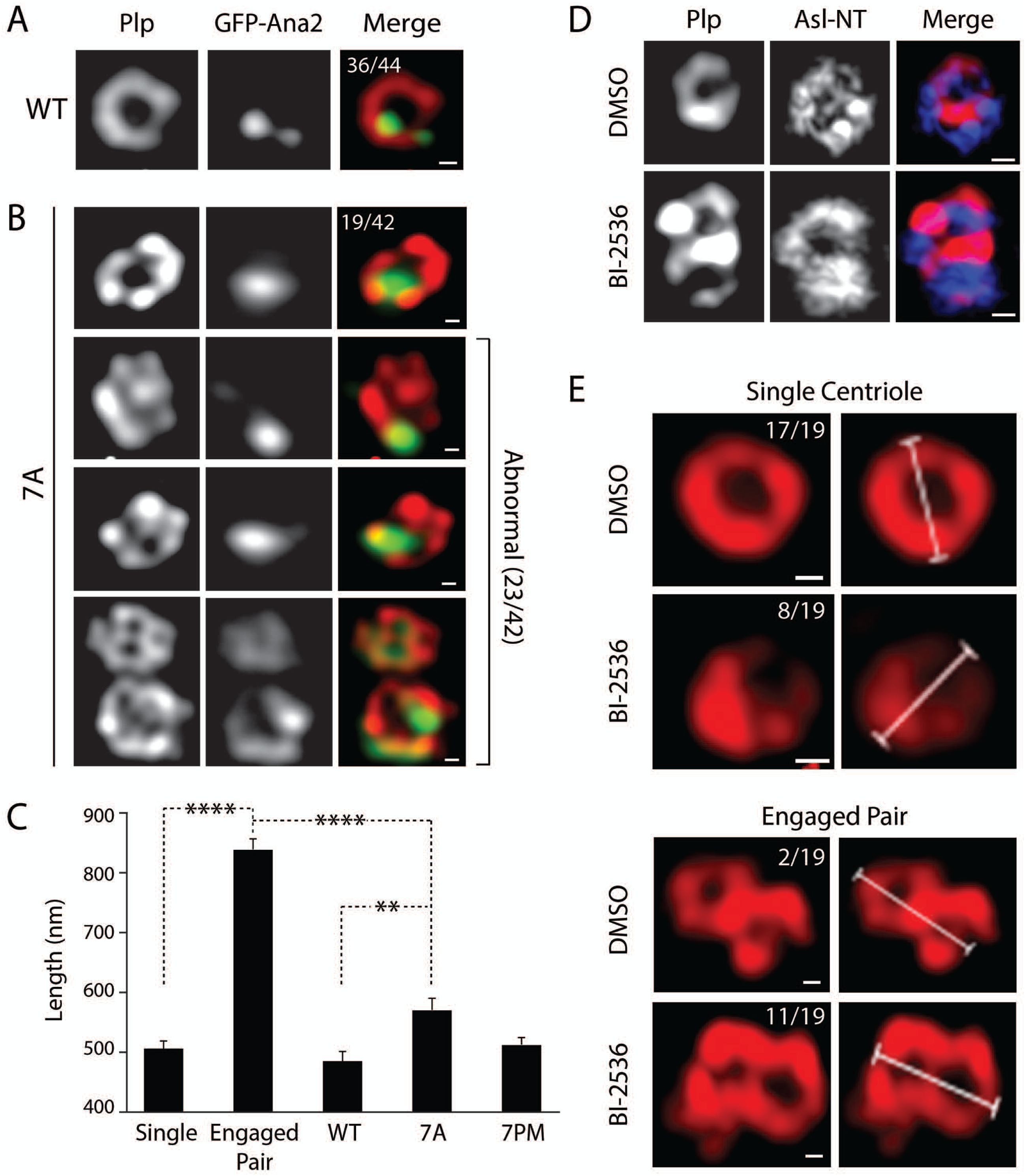
Centrioles containing Ana2-7A display an abnormal architecture but this is not due to a failure in disengagement. (A, B) The morphology of the centriole outer surface protein, PLP, is abnormal in Ana2-7A-expressing cells. Interphase centrioles in inducible GFP-Ana2-WT (A) or GFP-Ana2-7A (B) stable cell-lines were imaged using super-resolution microscopy (SR-SIM). Expression of GFP-Ana2 (green) was induced for 5 days. Cells were immunostained for PLP (red) and Hoechst-stained for DNA (not shown). The number of centrioles with normal (A and B, row 1) or abnormal (B, rows 2–4) morphology is indicated. Scale bars, 200 nm. (C) Graph shows average lengths of the major axis of centrioles in interphase cells. Measurements are shown for disengaged single centrioles in DMSO-treated cells (n=17), engaged centriole pairs in BI2536-treated cells (n=9), and centrioles in stable GFP-Ana2-expressing cells (WT, n=19; 7A, n=13; 7PM, n=34). Asterisks mark significant differences between treatments. Error bars, SEM. (D) Chemical inhibition of Polo blocks centriole disengagement in S2 cells. Cells were treated for 48 hours with either DMSO or the BI-2536 polo inhibitor, and interphase centrioles were imaged using (SR-SIM). Cells were immunostained for PLP (red) and Asl-NT (blue). Scale bars, 200 nm. (E) Examples of centriole measurements in control and Polo-inhibited interphase S2 cells. Cells were treated and prepared for SR-SIM as in D. The major axis of single centrioles and engaged pairs was measured. The numbers of measured single centrioles (upper panel) and engaged pairs (lower panel) are indicated. Scale bars, 200 nm.

One potential explanation for the abnormal PLP pattern within centriole centers is that the centrioles we imaged were not single centrioles, but rather two centrioles in a tightly engaged configuration. In other words, Ana2-7A expression may block mother-daughter centriole disengagement and because disengagement of the mother-daughter pair is a prerequisite for duplication (Tsou and Stearns, 2006), then this would provide a mechanistic explanation for Ana2-7A’s inhibition of centriole duplication. To test this hypothesis, we generated engaged centriole pairs in S2 cells by using the drug BI2536 to chemically inhibit Polo kinase (Fig. 5 D) whose activity is required for centriole separation in spermatocytes (Riparbelli et al., 2014). After 48 hours of drug treatment, most centrioles (58%, 11 of 19) in interphase cells remained engaged, as compared to 11% (2 of 19) in DMSO-control cells (Fig. 5 E). Not surprisingly, the average length of the major axis of an engaged pair (840 nm) was significantly greater than a single centriole (506 nm) (Fig. 5 C). The major axis length of an engaged pair was also significantly longer than centrioles in Ana2-7A expressing cells (Fig. 5 C), suggesting that the PLP-labelled objects in 7A mutant expressing cells are unlikely to be products of failed centriole disengagement. Instead, our findings support the conclusion that centrioles containing Ana2-7A are likely structurally abnormal and, to some extent, compromise both Ana2 recruitment to procentrioles and centriole duplication. If true, then a phosphorylated Ana2 N-terminus is an important structural constituent of centrioles and necessary to recruit Ana2 to the procentriole.

### Plk4 phosphorylates Ana2 in an ordered pattern

Ana2 binds and activates Plk4 and, in turn, is phosphorylated. The fact that multiple domains of Ana2 are modified raises the intriguing possibility that the phosphorylations occur sequentially, perhaps causing Ana2’s multiple activities to switch states (or levels) in an ordered succession. For example, our findings with Ana2-7PM suggest that if Plk4 first phosphorylated residues upstream of the STAN domain, then it would release from Ana2 and fail to phosphorylate the STAN domain, which seems unlikely because STAN phosphorylation is required to recruit Sas6 (Dzhindzhev et al., 2014; Ohta et al., 2014; Kratz et al., 2015; Moyer et al., 2015). To test this, we sampled a mixture of purified Ana2, Plk4 and MgATP at different time points and then mapped phospho-residues in Ana2 by MS/MS (Fig. 6 A). Strikingly, we found that phosphorylation was rapid in our conditions; at the earliest time point (<1 min), Ana2 was phosphorylated on N-terminal residues T33 and S38 as well as S318 within the STAN domain. At 1 and 5 min, Plk4 phosphorylated additional STAN domain residues. Phosphorylation of residues within the central region of Ana2 (S63/T69/S150/T159/T242) was first detected at 5 minutes and continued to accumulate until the last time point. Although the phosphorylation pattern of Ana2 may differ in cells, these in vitro findings suggest that Plk4 phosphorylates Ana2 residues in a specific order: 1) N-terminus (T33/S38), 2) C-terminus (STAN), and 3) central region.

**Figure 6.**
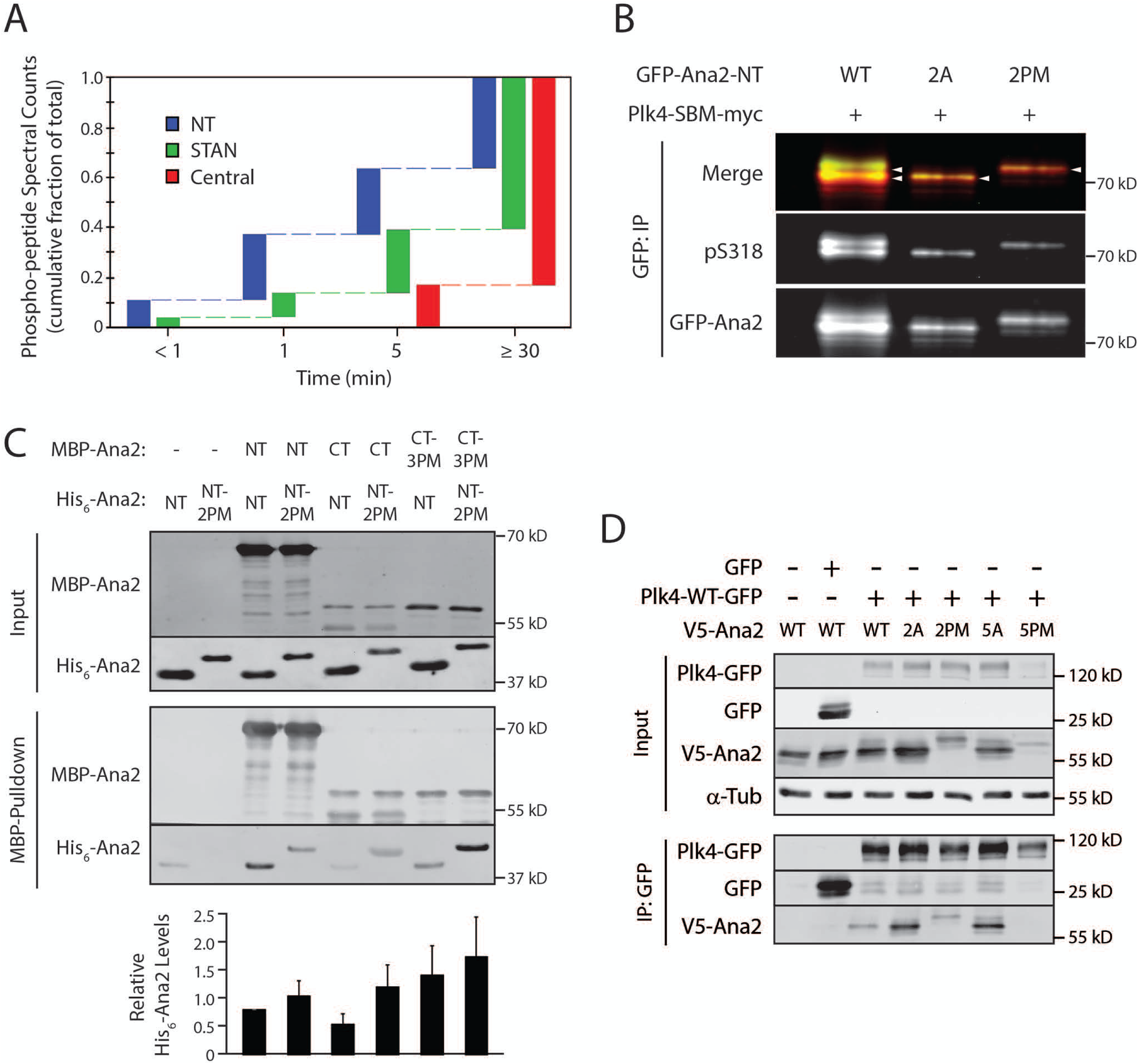
Plk4 phosphorylates Ana2 in an ordered pattern to regulate STAN domain activation and release Plk4. (A) Plk4 phosphorylates a specific pattern of Ana2 residues over time. Samples were taken at the indicated times from in vitro assays containing purified Plk4 1-317, GST-Ana2 and MgATP, and the phosphorylated residues of the sampled Ana2 identified by MS/MS. Using data pooled from four analyses, the spectral counts of peptides containing phosphorylated residues within the NT (containing T33, S38), central (S63, T69, S150, T159, T242), or STAN (S318, S370, S373, S395) regions were examined. The fraction of the maximal spectral count number seen at each time point is plotted. (Maximum spectral counts were observed in the last time point in every assay.) Note that the fraction measured at the earliest time point is highest (0.11) for NT residues and lowest (0) for central region residues. (B) Phosphorylation of STAN domain residue S318 is reduced by Ana2-2A but not by the Ana2-2PM mutant. Cells were depleted of endogenous Ana2 by RNAi for 7 days. On day 5, cells were co-transfected with the indicated GFP-Ana2 construct and active Plk4-SBM-myc, and the next day induced to express for 24 hours. Anti-GFP IPs were then prepared from lysates, and blots of the IPs probed for pS318 and GFP. A merge of GFP-Ana2 and pS318 blots is shown. Graph shows relative pS318 intensities in the IPs. For each treatment, intensities of pS318 were measured, normalized to GFP-Ana2 intensity levels, and then plotted relative to control (lane 1). (C) Phosphorylation increases the interaction between Ana2-NT and CT. The indicated combinations of MBP-or His_6_-tagged Ana2 NT (aa 1-317) or CT (aa 318–420) proteins were mixed for 30 min, after which the MBP construct was recovered by pull-down with amylose resin. Constructs were either wild-type or mutated with phosphomimetic (PM) substitutions: 2 residues (T33/S38) in NT (2PM), 3 residues (S318/S370/S373) in CT (3PM). Western blots of the washed resins were probed with anti-Ana2 antibody. Graph shows His_6_-Ana2 levels normalized to control pulldowns lacking MBP-Ana2. (D) Mutations in Ana2-NT (2A or 2PM) and central domain 5A do not prevent binding to Plk4 and, in fact, display stronger association with phospho-null substitutions (2A and 5A). In contrast, mutation of the central residues to phosphomimetic residues (5PM) eliminates Plk4 association. S2 cells were depleted of endogenous Ana2 by RNAi for 7 days; V5-Ana2 constructs were co-transfected with Plk4-GFP on day 5; cells were induced on day 6 to express the transgenes for 24 hours. Anti-GFP IPs were then prepared from lysates, and Western blots of the inputs and IPs probed for GFP, V5, and α-tubulin.

### Whereas phosphorylation of the NT residues T33 and S38 promotes Ana2 folding and hyperphosphorylation, modification of Ana2 central residues disrupts Plk4 association

The identification of T33 and S38 as early phosphorylation targets of Plk4 prompted us to reexamine the effects of Ana2 mutants on centriole duplication. We generated Ana2 mutants with single non-phosphorylatable alanine substitutions and transfected these into cells depleted of endogenous Ana2. As before, Ana2 depletion caused significant centriole loss which was rescued by replacement with Ana2-WT expression (Fig. S4 A). Interestingly, expression of only Ana2-S38A failed to rescue centriole numbers. Although S38 is near the Sas4-binding domain, both S38 mutants (S38A and S38D) bound Sas4 (when co-IPed from transfected cell lysates) (Fig. S4 B), and both also bound Plk4 (Fig. S4 C). Thus, expression of the S38A mutant inhibits centriole duplication, but this is likely not due to a defect in either Sas4 or Plk4 binding, unlike Ana2-7PM.

To understand how phosphorylation of the N-terminus contributes to Ana2 function, we next quantified STAN domain activation in phospho-mutants (alanine [A] and phosphomimetic [PM]) that cluster within the N-terminus (NT) (T33/S38; 2A and 2PM). Endogenous Ana2-depleted cells were co-transfected with GFP-Ana2 (WT or mutant) and catalytically-active Plk4-SBM. We then measured pS318 levels in GFP-Ana2 immunoprecipitates. As described previously, pS318 antibody reacted with Ana2-WT from cells co-expressing Plk4-SBM (Fig. 6 B, lane 1). Strikingly, detection of pS318 was confined to a single fast migrating band on SDS-PAGE, suggesting that expressed Ana2 cannot be hyper-phosphorylated when T33/S38 are mutated to alanines (Fig. 6 B, lane 2). In contrast, expressed Ana2-2PM migrated entirely as a slower migrating band on SDS-PAGE, suggesting that phosphomimetic Ana2 NT-2PM is completely hyper-phosphorylated (Fig. 6 B, lane 3). CD analysis of purified Ana2-2A and 2PM proteins revealed no difference in their spectra as compared to Ana2-WT (Fig. S3 D, E), suggesting that these mutations do not alter the structural integrity of Ana2. These findings suggest that modification of the N-terminus is required for maximal Ana2 phosphorylation. Although pS318 is detected in the Ana2-2A mutant (similar to the 7A mutant), further phosphorylation of STAN domain residues may be absent in Ana2-2A, possibly explaining the failure of Ana2-S38A to rescue centriole duplication.

Recently, a super-resolution microscopy study demonstrated that the N- and C-termini of Ana2 are positioned close to one another within the outer cartwheel of centrioles (Gartenmann et al., 2017), suggesting that Ana2 adopts a folded conformation. Therefore, we examined whether the N- and C-termini of Ana2 interact and, if so, whether this could be regulated by phosphorylation. We first examined in vitro binding of purified Ana2-NT (amino acids 1-317) and Ana2-CT (318-420) (containing mostly STAN domain) using pulldowns of MBP and His_6_-tagged proteins. As expected, Ana2-NT interacted with itself, as both constructs contain the CC (Fig. 6 C); this interaction was not altered in Ana2-NT containing phosphomimetic T33E/S38D (2PM) substitutions, suggesting that phosphorylation of these residues is not important for oligomerization through the CC. Notably, Ana2-NT and CT do not bind (Fig. 6 C), and we confirmed this an inconsistent/weak interaction by Y2H (Fig. S4 D). Next, we examined binding between Ana2-NT-2PM and Ana2-CT containing PM substitutions of its three most conserved phosphorylated STAN residues (S318/S370/S373). Surprisingly, mutation of the NT sites to PM residues did not interfere with the NT-CT interaction. In fact, when Ana2 pulldown bands were quantified (Fig. 6 C, graph), the opposite is found: PM residues in either the NT or CT construct increase the average amount of interaction by 2-3 fold. Interestingly, mutation to phosphomimetic of both domains, induces the highest interaction (~3.5 fold), suggesting that phosphorylation of Ana2 increases the interaction between its N- and C-termini.

Next, we examined the effects of phosphorylation of the NT (2A or 2PM) and the Plk4-targeted residues of Ana2’s central region (S63/T69/S150/T159/T242; 5A or 5PM) on Plk4 association by co-IP. When V5-Ana2 (WT or mutant) was co-transfected with Plk4-GFP into cells, non-phosphorylatable Ana2-2A and 5A mutants interacted strongly with Plk4 (Fig. 6 D, lanes 4, 6), whereas WT and phosphomimetic 2PM interacted weakly with Plk4 (Fig. 6 D, lanes 3, 5). Strikingly, Ana2-5PM failed to interact with Plk4 (Fig. 6 D, lane 7), suggesting that phosphorylation of the central region causes Ana2 to dissociate from Plk4. Consistent with its inability to bind Plk4, expression of Ana2-5PM, but not Ana2-5A, failed to rescue centriole duplication in cells depleted of endogenous Ana2 (Fig. S4 E). Taken together, our data support the model that phosphorylation of NT residues T33/S38 and CT residues S318/S370/S373 promote Ana2 folding. Subsequently, phosphorylation of the 5 central residues causes Plk4 dissociation.

## Discussion

Our investigation of Plk4 and Ana2 reveals a complex interaction: they cooperate to relieve autoinhibition and promote centriole assembly through a series of phosphorylation events (Fig. 7). The activities of Polo-like kinase family members are initially restrained by multiple mechanisms, including autoinhibition (Lowery et al., 2005). We demonstrated previously that Plk4 is no exception and uses a linker (L1) adjacent to the kinase domain to autoinhibit (Klebba et al., 2015a) (Fig. 7 A), similar to Plk1 (Xu et al., 2013). We also showed that relief of autoinhibition requires Polo-box 3 (PB3) as well as an unidentified extrinsic factor (Klebba et al., 2015a). STIL is likely the unidentified factor because it binds PB3 and stimulates Plk4 activity in cells (Moyer et al., 2015; Arquint et al., 2015). Additionally, Arquint et al. (2015) identified a second STIL-binding site in Plk4, positioned somewhere between the kinase domain and Polo Box 1 (PB1). Using a combination of immunoprecipitation, yeast two-hybrid analysis, and in vitro protein binding methods, we show that *Drosophila* Plk4 also contains a second Ana2-binding site that maps to a conserved, leucine-rich segment (34 amino acids) located immediately downstream of the kinase domain and is predicted to form a coiled-coil (CC). The crystal structure of the STIL-CC with the single α-helix within PB3 reveals a leucine zipper-type CC interaction (Arquint et al., 2015). We predict that a similar structure forms between Ana2-CC and the newly identified putative CC region of Plk4. Intriguingly, the corresponding region of human Plk4 possessesα-helical content (Wong et al., 2015), suggesting that this interaction site could be conserved between human Plk4 and STIL. However, we note that there are known differences in the quaternary structures of STIL and Ana2 with Plk4. For example, whereas the STIL-CC is monomeric when complexed with human PB3, Ana2-CC forms a tetramer (Cottee et al., 2015) and also has been shown to simultaneously bind eight dynein light chain (LC8) subunits (Slevin et al., 2014). Given the significance of their interaction, an important goal of future studies is to determine how the Ana2-CC associates with both PB3 and the putative CC in Plk4 and to also determine the stoichiometry of this complex.

**Figure 7.**
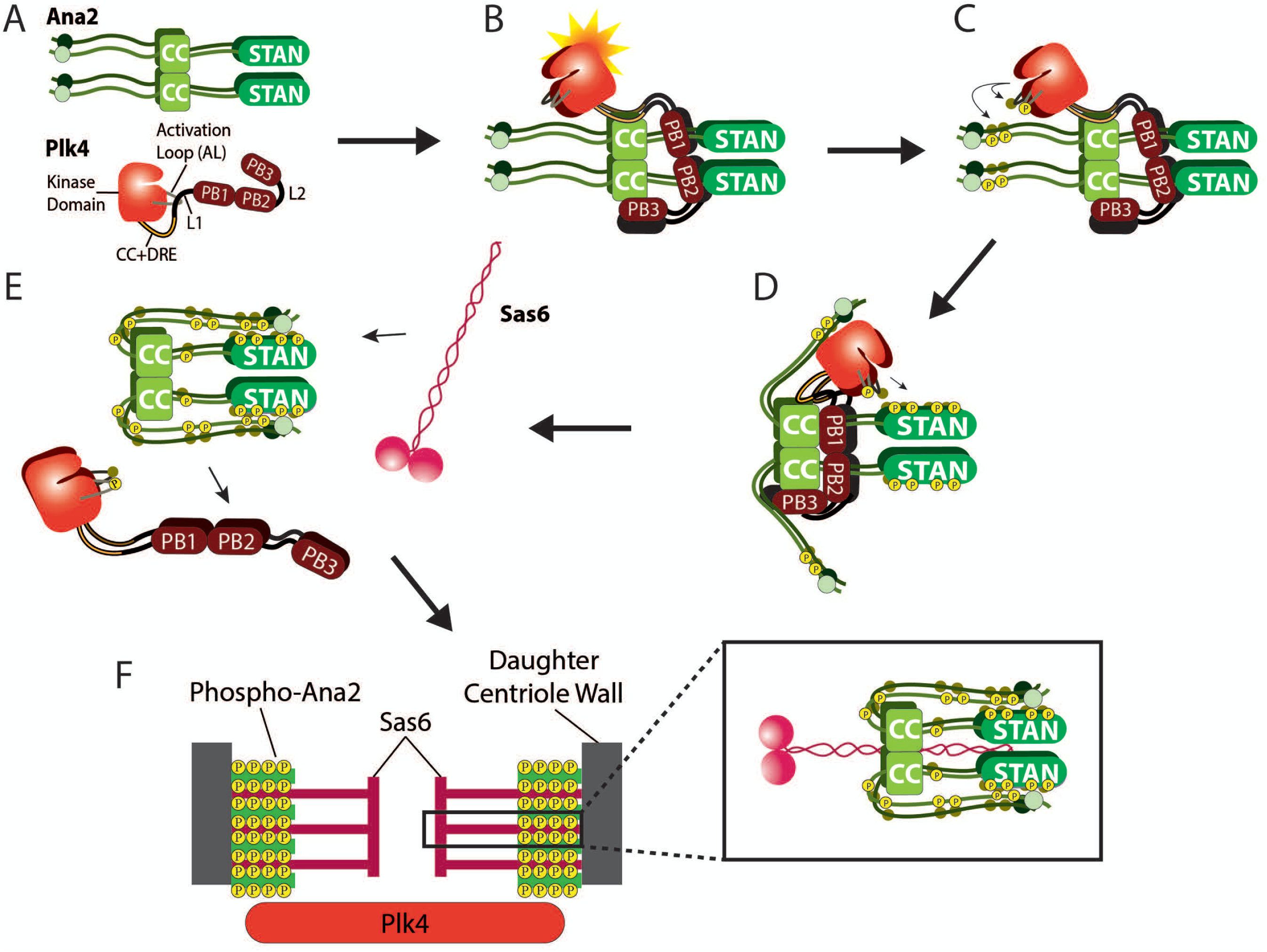
Proposed model depicting the mechanism of Plk4 and Ana2 interdependent activation to achieve centriole assembly. (A) Plk4 and Ana2 exist initially in nonfunctional states. Nascent monomeric Plk4 is inactive, a state maintained by L1 which inhibits the phosphorylation of the activation loop needed for full enzymatic activity. Similarly, Ana2 is initially nonfunctional, i.e., does not support centriole assembly until phosphorylated by Plk4 and exists an ‘open’ conformation. (B) Ana2 targets centrioles where it binds Plk4. Centriole targeting of Ana2 occurs through an unidentified mechanism that requires Plk4 kinase activity. The CC domains in tetrameric Ana2 bind two sites in Plk4 (PB3 and the putative CC adjacent to the kinase domain). Consequently, (1) L1-mediated Plk4 autoinhibition is relieved, and (2) kinase domains are clustered and oriented, optimizing their *trans*-autophosphorylation in the Plk4 homodimeric state. (C) Disinhibited Plk4 autophosphorylates its activation loops and phosphorylates Ana2 on two NT residues, T33 and S38. (D) NT phosphorylation induces a conformational change of Ana2 that enhances Plk4 phosphorylation of the STAN domain, which is known to recruit Sas6 to the procentriole assembly site. (E) Ana2’s conformational change exposes five residues within the central region of Ana2 to Plk4, and subsequently their phosphorylation causes Ana2 to release from Plk4. (F) Ana2 tetramers incorporate into the growing procentriole to stabilize the vertical stacking of Sas6 cartwheels (as proposed by Cottee et al., 2015). The hyper-phosphorylated Ana2-NT may also promote additional stabilizing interactions with proteins positioned near centriolar microtubules. This is important in order to maintain structural integrity of the organelle and, if compromised, could interfere with procentriole assembly.

Plk4 and Ana2 binding has consequences for centriole duplication, not because this binding is required to target Ana2 to the site of procentriole assembly, but because binding appears to initiate changes in the functions or activities of both proteins. For example, our findings suggest that Ana2 relieves Plk4 autoinhibition by a mechanism requiring PB3, which is consistent with our previous work (Klebba et al., 2015a). However, Ana2 will even activate a minimal Plk4 construct (containing the kinase domain and predicted CC) that lacks the autoinhibitory L1 domain. Therefore, we propose that when Ana2 complexes with multiple Plk4 proteins (presumably through the putative Plk4-CC), the Plk4 kinase domains become clustered in orientations that facilitate *trans*-autophosphorylation of their activation loops (Fig. 7 B) (see also Lopes et al., 2015). Although disinhibition is an important step in unlocking Plk4 kinase activity, we propose that the release of autoinhibition may not, by itself, trigger full Plk4 activation. Instead, the catalytic activity of Plk4 is further stimulated by a distinct step of clustering and orienting kinase domains that optimizes *trans*-autophosphorylation.

Though Ana2/STIL is the first described stimulator of Plk4 activity, other unidentified activators probably exist. For example, work in *C. elegans* embryos has shown that Plk4/ZYG1 is recruited to centrioles before SAS-5 (the worm homolog of Ana2/STIL) (Pelletier et al., 2006; Delattre et al., 2006). So what activates Plk4 before Ana2’s arrival? *Drosophila* Asterless (Asl) is an interesting candidate because it targets Plk4 to centrioles and plays additional roles in stabilizing the kinase (Dzhindzhev et al., 2010; Klebba et al., 2015b). Identifying other Plk4 activators will be essential to determine how Ana2 targets centrioles because, intriguingly, Plk4 kinase activity is required for STIL’s centriolar recruitment (Moyer et al., 2015; Zitouni et al., 2016). Although it has been proposed that direct binding to Plk4 is responsible for STIL recruitment (Ohta et al., 2015; Moyer et al., 2015), we feel this is an open question. Previous work has shown that the Ana2-CC is required for its centriole localization and Plk4 binding (Ohta et al, 2014; Arquint et al., 2015; Moyer et al., 2015) and this observation forms the basis for the current model of Ana2’s centriole targeting. But we found that CC deletion is remarkably pleiotropic, because it also disrupts oligomerization and Sas4 binding. [Possibly, failure to oligomerize compromises most Ana2 interactions, including with Cdk1-cyclinB (Zitouni et al., 2016).] Our work demonstrates that Ana2-7PM fails to bind Plk4 but still successfully localizes to the procentriole. Therefore, we propose that direct binding to Plk4 is not required for Ana2 localization. Possibly another centriole surface protein(s) can, after phosphorylation by Plk4, function as a high-affinity binding site for Ana2/STIL. Additional work is needed to solve this problem.

Ana2 stimulates Plk4 activity and, in turn, is extensively phosphorylated along its length ((Dzhindzhev et al., 2014; Ohta et al., 2014; Kratz et al., 2015). Our data suggest that Plk4 likely initially targets N-terminal residues T33/S38, which then promote STAN domain phosphorylation. However, our data are also compatible with a model where residues of both NT and STAN regions are phosphorylated contemporaneously, with the end result being the same. We propose that Ana2 exists in an open conformation, where the NT and STAN domain are separated (Fig. 7 A). In this model, phosphorylation of T33/S38 induces Ana2 to fold, and the conformational change may be a functionally important event -- it could, for example, facilitate Plk4’s further phosphorylation of key residues in the STAN domain that promote the subsequent recruitment of Sas6 to the procentriole assembly site (Fig. 7 C, D). This model predicts our observation that, when expressed in cells, the non-phosphorylatable Ana2-NT-2A mutant (T33A/S38A) does not display the mobility-shifted hyper-phosphorylated band seen with Ana2 WT or 2PM expression (Fig. 6 B). Our interpretation of this result is that NT phosphorylation is a prerequisite for the complete modification of all the Plk4-targeted residues of Ana2. Further supporting this model is our finding that the Ana2 S38A single point mutant completely fails to rescue centriole duplication in cells depleted of endogenous Ana2 (Fig. S4 A). If S38 phosphorylation is a prerequisite for the full course of Plk4-mediated phosphorylation, then failure of centriole duplication is expected for the S38A mutant. Similarly, the 7A mutant is also not hyperphosphorylated, likely due to the mutation of the two NT residues (Fig. 4 D). If Ana2 initially adopts an ‘open’ conformation, its extensive phosphorylation could drive a conformational change that promotes Sas6 binding, thus enabling procentriole assembly.

Expression of phopsho-Ana2 mutants (7A or 7PM) had an unexpected effect; neither mutant rescued the loss of centrioles phenotype in endogenous Ana2-depleted cells. However, this phenotype is reached as a result of disrupting different Ana2-dependent steps in centriole assembly. Our results suggest that, after STAN domain modification, Plk4 then targets several residues within the central region of Ana2, generating a hyperphosphorylated form of Ana2 that dissociates from Plk4. Extensive phosphorylation of the central region appears to inhibit Plk4 binding, as observed with the Ana2-5PM and 7PM mutants.

Ours is not the first description of a mechanism that regulates the Plk4-STIL/Ana2 interaction. During mitosis, Cdk1-cyclinB binds the CC domain in STIL and prevents Plk4 association (Zitouni et al., 2016). This ensures that Plk4 activation and STAN phosphorylation are restricted to periods of the cell cycle when Cdk1 activity is low, such as during mitotic exit when Ana2 and Sas6 are first recruited to centrioles (Dzhindzhev et al., 2014). This same period is when the Ana2/Plk4 interaction that we describe probably occurs.

Lastly, we propose that hyper-phosphorylated Ana2 releases Plk4 and incorporates into the nascent procentriole where it makes contact with Sas6 and other centriole proteins (Fig. 7 F). Phosphorylation of Ana2/STIL may regulate its ability to organize Sas4/CPAP into the horizontal struts that connect the centriole spokes with microtubules (Hatzopoulos et al., 2013). Notably, STIL/SAS-5 turnover is rapid between centrioles and cytoplasm (Vulprecht et al., 2012; Delattre et al., 2004), and may act as a transporter for centriolar components (Hatzopoulos et al., 2013) - - yet another function that may be regulated by phosphorylation. Thus, we interpret centriole loss in cells expressing Ana2-7A to be a result of defective Ana2 recruitment to the procentriole. It will be instructive to determine the path of recruitment to the procentriole, whether Ana2 is first recruited to the lumen of the mother as demonstrated for Sas6 (Fong et al., 2014), or instead is directly recruited to the procentriole. Moreover, we recently reported that Ana2 interacts with multiple centriole proteins in a yeast-two hybrid screen of centrosome components, including Asl, Cep135 and Ana1 (Galletta et al., 2016). Whether Plk4 activity alters these interactions remains to be determined. In any case, it is increasingly clear that Plk4 is a fundamentally important regulator of the interaction landscape of centriolar proteins that is needed for the seemingly complex molecular choreography that underlies centriole assembly.

## Materials and methods

### Cell culture and double-stranded RNAi

*Drosophila* S2 cell culture, in vitro dsRNA synthesis, and RNAi treatments were performed as previously described (Rogers and Rogers, 2008). Cells were cultured in Sf900II SFM media (Life Technologies). RNAi was performed in 6 well plates. For immune precipitation assays, cells (40-90% confluency) were treated with 10 μg of dsRNA in 1ml of media and replenished with fresh media/dsRNA every day for 7 days. For immunofluorescence microscopy, cells were transfected with 40 μg dsRNA on days 0, 4 and 8. A ~550 bp control dsRNA was synthesized from DNA template amplified from a non-coding sequence of the pET28a vector (Clontech) using the primers 5’-ATCAGGCGCTCT TCCGC and 5’-GTTCGTGCACACAGCCC. (All primers used for dsRNA synthesis begin with the T7 promoter sequence 5’-TAATACGACTCACTATAGGG, followed by template-specific sequence). dsRNA targeting the Ana2 UTR was synthesized from EST LD22033 template by first deleting the Ana2 cDNA, and then joining 91 bp of 5’UTR with 78 bp of 3’UTR to create the following sequence: 5’-AGTTCCACCCCTAAGTCGATCGACTTCCAATTGGACAGATTCTCCCGCTCGAATTTAATTTAATCGGCAAATATAAACAAATACGCTCCAAAAGCATG TACAATGTTCGTTTTGTTATTTATGCATATGTCTATTTGCGATTTAAGTGGAAATATATTTCAATACACGG-3’. This template was amplified using the primers 5’-CAGATTCTCCC GCTCG and 5’-TTCCGTGTATTGAAATATATTTCC. Immunoblotting confirmed that Ana2 UTR RNAi depleted endogenous Ana2 by ~80-90% (Fig. S1 C).

### Immunoblotting

S2 cell extracts were produced by lysing cells in cold PBS and 0.1% Triton X-100. Laemmli sample buffer was then added and samples boiled for 5 min. Samples of equal total protein were resolved by SDS-PAGE, blotted, probed with primary and secondary antibodies, and scanned on an Odyssey imager (Li-Cor Biosciences). Care was taken to avoid saturating the scans of blots. Antibodies used for Western blotting include rabbit anti-Ana2 (our laboratory), rat anti-Cep135 (our laboratory), rat anti-Asl-A (amino acids 1-374) (our laboratory), mouse anti-V5 monoclonal (Life Technologies), mouse anti-GFP monoclonal JL8 (Clontech), mouse anti-myc (Cell Signaling Technologies), mouse anti-α tubulin monoclonal DM1A (Sigma-Aldrich), mouse anti-GST (Cell Signaling), and mouse anti-FLAG monoclonal (Sigma-Aldrich) used at dilutions ranging from 1:1,000-30,000. IRDye 800CW secondary antibodies (Li-Cor Biosciences) were prepared according to the manufacturer’s instructions and used at 1:3,000 dilutions.

To generate the anti-phospho-specific T172 Plk4 antibody, rabbit polyclonal antibodies were raised against the following phospho-peptide: Acetyl-PDERHM(pT)MCGTPN. A non-phospho-peptide (Acetyl-PDERHMTMCGTPN) was also synthesized (ThermoFisher). To generate the anti-phospho-specific pS318 Ana2 antibody, rat polyclonal antibodies were raised against the following phosphor-peptide: AKPNTEK{pSer}MVMNELAC. A non-phosphopeptide (AKPNTEKSMVMNELAC) was also synthesized (ThermoFisher). Antibodies were affinity-purified from antisera using Affi-Gel 10/15 resin (BioRad Laboratories) coupled to the phosphopeptide. The eluted material was then pre-absorbed over an Affi-Gel column coupled to the non-phosphospecific peptide. The eluate was collected and concentrated using 10K Ultrafree 2 ml concentrators (Millipore). Antibodies were used at a 1:1000 dilution.

### Mass Spectrometry

Samples for MS/MS analysis were first resolved by SDS-PAGE and Coomassie stained. Bands of interest were cut from gels and then processed for MS. Gel pieces containing Ana2 were reduced (10 μM dithiothreitol, 55°C, 1hr), alkylated (55 mM iodoacetamide, room temperature, 45 min), and trypsin digested (~1 μg trypsin, 37°C, 12 hrs; for one experiment, a sequential chymotrypsin/trypsin digest was used) in-gel, and then extracted. Peptide samples were loaded onto a Zorbax C_18_ trap column (Agilent Tech., Santa Clara, CA) to desalt the peptide mixture using an on-line Eksigent (Dublin, CA) nano-LC ultra HPLC system. The peptides were then separated on a 10 cm Picofrit Biobasic C_18_ analytical column (New Objective, Woburn, MA). Peptides were eluted over a 90 min linear gradient of 5-35% acetonitrile/water containing 0.1% formic acid at a flow rate of 250 nL/min, ionized by electrospray ionization (ESI) in positive mode, and analyzed on a LTQ Orbitrap Velos (Thermo Electron Corp., San Jose, CA) mass spectrometer. All LC MS analyses were carried out in “data-dependent” mode in which the top 6 most intense precursor ions detected in the MS1 precursor scan (m/z 300-2000) were selected for fragmentation via collision induced dissociation (CID). Precursor ions were measured in the Orbitrap at a resolution of 60,000 (m/z 400) and all fragment ions were measured in the ion trap.

MS/MS data used for Fig. 3A was accumulated from 5 experiments and generated by two facilities [Taplin Mass Spectrometry Facility (Harvard Medical School) and the NHLBI Proteomics Core Facility (NIH)]. Typically, coverage of Ana2 was ~85%; coverage of residues 13-65 was often problematic. For Fig. 6A, coverage ranged from ~85-95%; residues 1-12, 414-420 were never recovered. For Fig. S3 C, best coverage was ~80-85%; coverage of residues 13-65 was often problematic.

### Immunofluorescence microscopy

S2 cells were fixed and processed as previously described (Rogers and Rogers, 2008) by spreading S2 cells on concanavalin A–coated glass-bottom dishes and fixing with ice-cold methanol. Primary antibodies were diluted to concentrations ranging from 1 to 20 μg/ml. They included rabbit anti-PLP (Rogers et al., 2009), guinea pig anti-Asl (Klebba et al., 2013), rat anti-Cep135, rat anti-Asl-NT, and mouse anti-V5 (Life Technologies) antibodies. Goat secondary antibodies (conjugated with Cy2, Rhodamine red-X, or Cy5 [Jackson ImmunoResearch]) were used at 1:1,500. Hoechst 33342 (Life Technologies) was used at a final concentration of 3.2 μM. Cells were mounted in 0.1 M *n*-propyl galate, 90% (by volume) glycerol, and 10% PBS solution. Specimens were imaged using a DeltaVision Core deconvolution system (Applied Precision) equipped with an Olympus IX71 microscope, a 100× objective (NA 1.4), and a cooled charge-coupled device camera (CoolSNAP HQ2; Photometrics). Images were acquired with softWoRx v1.2 software (Applied Science). Super resolution microscopy was performed using a Zeiss ELYRA S1 (SR-SIM) microscope equipped with an AXIO Observer Z1 inverted microscope stand with transmitted (HAL), UV (HBO) and solid-state (405/488/561 nm) laser illumination sources, a 10× objective (NA 1.4), and EM-CCD camera (Andor iXon). Images were acquired with ZEN 2011 software.

### Constructs and transfection

Full-length cDNAs of *Drosophila* Ana2, Plk4, Sas4, and Sas6 were subcloned into a pMT vector containing in-frame coding sequences for EGFP, V5, or myc under control of the inducible metallothionein promoter. PCR-based site-directed mutagenesis with Phusion polymerase (ThermoFisher) was used to generate the various Ana2 and Plk4 deletion and point mutants. Transient transfections of S2 cells were performed as described (Nye et al., 2014). Briefly, ~2–5 × 10^6^ cells were pelleted by centrifugation, resuspended in 100 μl of transfection solution (5 mM KCl, 15 mM MgCl_2_, 120 mM sodium phosphate, 50 mM D-mannitol, pH 7.2) containing 1-3 μg of purified plasmid, transferred to a cuvette (2-mm gap size), and then electroporated using a Nucleofector 2b (Lonza), program G-030. Transfected cells were diluted immediately with 0.5 ml of SF-900 II medium and plated in a six-well cell-culture plate. Typically, cells were allowed 24 hrs to recover before further manipulation. Expression of all constructs was induced by addition of 100 μM–1 mM copper sulfate to the culture medium.

### Yeast Two-Hybrid assay

Yeast two-hybrid (Y2H) experiments were carried out using the Matchmaker Gold Y2H system (Clontech) with significant modifications. pDEST-GADT7 and pDEST-GBKT7, modified versions of Matchmaker vectors compatible with the Gateway cloning system (Life Technologies), were used (Rossignol et al., 2007). pDEST-GADT7 and pDEST-GBKT7 contain the 2 μ and pUC ori for growth in yeast and bacteria. Both contain the Gateway cassette and utilize the ADH1 promoter to drive expression in *Saccharomyces cerevisiae*. pDEST-GADT7 fuses the SV40 nuclear localization signal, the GAL4 activation domain and the HA epitope tag to the amino-terminus of the protein encoded by DNA inserted into the Gateway cassette. pDEST-GBKT7 fuses the SV40 nuclear localization signal, the GAL4 DNA binding domain and the c-Myc epitope tag to the amino-terminus of the protein encoded by DNA inserted into the Gateway cassette. pDest-pGBKT7 was modified by yeast-mediated recombination to confer resistance to ampicillin instead of kanamycin. pDEST-GADT7 and pDEST-GBKT7 plasmids containing fragments encoding the protein regions to be tested for interaction were transformed into Y187 and Y2HGold yeast strains respectively using standard techniques. Liquid cultures of yeast carrying these plasmids were grown at 30°C, with shaking, to an OD600 ~0.5 in SD – leu or SD –trp media, as appropriate, to maintain plasmid selection. Interactions were tested by mating, mixing 20 μl each of a Y187 strain and a Y2HGold strain in 100μl of 2X YPD media in a 96 well plate. Mating cultures were grown for 20-24 hrs at 30°C with shaking. Cells were pinned onto SD –leu –trp (DDO) plates to select for diploids carrying both plasmids, using a Multi-Blot Replicator (VP 407AH, V&P Scientific) and grown for 5 days at 30°C. These plates were then replica plated onto DDO, SD – ade –leu –trp –ura (QDO), SD – leu –trp + Aureobasidin A (Clontech) + X-**α**-Gal (Clontech, Gold Biotechnology) (DDOXA) and/or SD – ade –leu –trp – ura + Aureobasidin A + X-**α**-Gal (QDOXA). Replica plates were grown for 5 days at 30°C. Interactions were scored based on growth and/or blue color, as appropriate.

### In vitro kinase assays

Bacterially-expressed constructs of *Drosophila* Plk4 (amino acids 1-317 or 1-602) C-terminally tagged with FLAG-His_6_, full-length His-MBP-Plk4 (gift from M. Bettencourt-Dias) and full-length *Drosophila* Ana2 N-terminally tagged with Glutathione S-Transferase (GST-Ana2) were purified on HisPur (ThermoFisher), amylose (NEB) or glutathione resin (NEB), respectively, according to manufacturers’ instructions. Prior to assay, Plk4 was pre-treated with λ-phosphatase (NEB) (because it autophosphorylates to some extent when expressed in bacteria). Samples of protein reagents were resolved by SDS-PAGE and scans of the Coomassie-stained gels analyzed by densitometry (ImageJ, NIH) to determine protein purity. Total protein concentrations were measured by Bradford assay (BioRad). The total protein and purity measurements were used to calculate the concentration of each protein reagent. (Contaminants and proteolytic fragments are excluded by this calculation.) In vitro phosphorylation assays were performed with the indicated proteins at different molarities (see Results) by incubation with 100 μM ATP for the indicated times at 25°C in reaction buffer RXN1 [40 mM Na HEPES (pH 7.3), 150 mM NaCl, 5 mM MgCl_2_, 1 mM DTT, 10% (by volume) glycerol]. Samples were resolved by SDS-PAGE, and proteins visualized by Coomassie staining. Phosphorylation of protein substrates was evaluated by including γ-^32^P-ATP in assays and, subsequently, the presence of radiolabeled substrates detected by autoradiography or phosphorimaging (STORM, GE Healthcare) of dried gels. Phosphorylated residues within proteins were identified by tandem mass spectrometry (Table S1) of purified bacterially-expressed proteins phosphorylated in vitro (described above) in the presence of non-radioactive ATP. For GST pull-down assays, purified proteins were concentrated using Amicon Ultra spin concentrators (EMD Millipore). GST or GST-Ana2 was immobilized on glutathione agarose (Pierce), mixed with His_6_-FLAG-Plk4 1-317, rocked at 25^o^C for 30 min, and pelleted at 500 *g* for 1min. Supernatants and washed pellets were analyzed by anti-FLAG and GST immunoblots.

### In vitro binding assays

Bacterially-expressed constructs of *Drosophila* Ana2 (amino acids 1-317 or 318-420) N-terminally tagged with His_6_ or maltose binding protein (MBP) were purified on HisPur (ThermoFisher), amylose (NEB), respectively, according to manufacturers’ instructions. MBP-Ana2 was not eluted from the resin for binding assays. Proteins were analyzed by SDS-PAGE and equimolar amounts of MBP-Ana2 and His_6_-Ana2 were mixed together in buffer RXN1. Samples were incubated for 30 minutes at room-temperature with shaking, followed by 8 washes with buffer RXN1. Resin was resuspended in sample buffer and subsequently run on SDS-PAGE for analysis by Western blotting.

### GFP immunoprecipitation assays

GFP-binding protein (GBP) (Rothbauer et al., 2008) was fused to the Fc domain of human IgG (pIg-Tail) (R&D Systems), tagged with His_6_ in pET28a (EMD Biosciences), expressed in *E. coli* and purified on HisPur resin (ThermoFisher) according to manufacturer’s instructions (Buster et al., 2013). Purified GBP was bound to magnetic Dyna Beads (ThermoFisher), and then crosslinked to the resin by incubating with 20mM dimethyl pimelimidate dihydrochloride in PBS, pH 8.3, 2 hours, 22°C, and then quenching the coupling reaction by incubating with 0.2 M ethanolamine, pH 8.3, 1 hour, 22°C. Antibody-coated beads were washed three times with PBS-Tween20 (0.02%), then equilibrated in 1.0 ml of cell lysis buffer (CLB; 50 mM Tris, pH 7.2, 125 mM NaCl, 2 mM DTT, 0.1% Triton X-100, 1x protease inhibitor cocktail [Roche] and 0.1 mM PMSF) or cell lysis buffer containing phosphatase inhibitors for phospho-specific antibody detection (CLB+; CLB with 200 mM NaF, 150 mM β-glycerophosphate, 1 mM Na3VO4). Transfected cells expressing recombinant proteins were lysed in CLB or CLB+, and the lysates clarified by centrifugation at 16,100 × g for 5 min, 4^o^C. 0.5–1% of the inputs were used for immunoblots. GBP-coated beads were rocked with lysate for 30 min, 4°C or 10 min, 22^o^C, washed four times with 1 ml CLB or CLB+, and then boiled in Laemmli sample buffer.

### Circular Dichroism (CD)

N-terminally GST-tagged Ana2 (WT or mutants) were bacterially expressed and then purified from clarified bacterial lysate using glutathione resin (Thermo Scientific) following manufacturer’s instructions. The GST tag was removed by incubating the resin-bound protein with PreScission Protease (GE Healthcare) overnight, 4°C. Protease-cleaved protein was recovered in the column flow-through, and the material was then passed through new glutathione resin to remove any remaining GST-tagged protease and uncut GST-tagged Ana2. Samples were analyzed by SDS-PAGE. Full length His_6_-Ana2 wild-type, 2A, 2PM, 7A, and 7PM constructs were diluted to 0.10 mg/mL in CD buffer (10 mM sodium phosphate, pH 7.4, 50 mM NaF). Spectra were acquired in a 1-mm-path length cuvette at 20°C from 260 to 185 nm with a step size of 0.5 nm every 1.25 seconds using a Chirascan-plus CD spectrometer (Applied Photophysics, Leatherhead, United Kingdom). A CD buffer spectrum was subtracted from each Ana2 spectrum and the resulting curves were smoothed using Chirascan-plus software. Representative traces (Fig. S3 E) are shown from data collected on two independent experimental days.

### Statistical analysis and curve fitting

Means of measurements were analyzed for significant differences by one-way ANOVA followed by Tukey’s post-test (to evaluate differences between treatment pairs) using Prism 7 (GraphPad) software. Means are taken to be significantly different if P < 0.05. P values shown for pairwise comparisons of Tukey’s post-test are adjusted for multiplicity. In figures, “*” indicates 0.05 > P ≥ 0.01, “**” indicates 0.01 > P ≥ 0.001, “***” indicates 0.001 > P ≥ 0.0001, “****” indicates 0.0001 > P, and “ns” indicates P ≥ 0.05 for the indicated pairwise comparison. Error bars in all figures indicate standard error of the mean (SEM).

Plots of in vitro phosphorylation assays were best-fit to rational functions (quadratic/quadratic) using nonlinear regression (MatLab, MathWorks). Initial reaction rates were obtained from the slopes of the plots covering the 0-30 min time points; differences in the slopes were analyzed for significance using Prism 7. In Fig. 2 C, the best-fit curves for assays that include Ana2 all have horizontal asymptotes (limits of Plk4 autophosphorylation) that do not differ significantly. In the case of the control (lacking Ana2), Plk4 activity is low and our measurements probably lie within the initial, exponential phase of the reaction profile, so this curve does not have a calculated limit. However, even assuming that the control reaction would always follow an exponential path, control Plk4 would require more than 7 hrs to reach a level of autophosphorylation similar to the calculated asymptotic limits of the Ana2-containing reactions.

The graph in Fig. S2 C was generated by first cutting appropriate Coomassie-stained bands from SDS-PAGE gels of samples from in vitro phosphorylation assays (e.g., Fig. S2 A), measuring radioactivity in a liquid scintillation counter (LS 6500, Beckman Coulter), and then best-fitting the data to third order polynomial curves using Prism 7.

## Online supplemental material

Fig. S1 shows specificity of anti-Ana2 antibodies, and that the Ana2-CC domain is necessary for Plk4 association. This figure shows that the putative Plk4-CC and PB3 domains interact with Ana2-CT by Y2H analysis and that the putative Plk4-CC is conserved. Fig. S2 shows that Plk4 protein containing L1 (Plk4-1-602) has greatly reduced kinase activity compared to a truncation mutant lacking L1 and is not activated by Ana2. This figure also demonstrates the specificity of the anti-pT172 antibody. Fig. S3 shows conservation of Ana2-NT phospho-residues and summarizes the in vivo Ana2 phosphorylation assay. It shows that Plk4 does not phosphorylate Sas6 in vitro, and that Ana2 phospho-mutants are not misfolded as determined by CD. It also shows that centrioles containing Ana2-7A recruit PCM during mitosis, and demonstrates the specificity of anti-pS318 Ana2, Cep135 and Asl-NT antibodies used in this study. Fig. S4 shows that Ana2 S38A and 5PM mutants fail to rescue centriole duplication, and that Ana2-S38A/PM does not prevent Sas4 or Plk4 binding. It also show that Ana2 N- and C-termini weakly interact via Y2H. Table S1 lists the Ana2 phospho-sites identified by tandem MS in this study.

## Acknowledgements

We thank P. Krieg for editing and M. Bettencourt-Dias for providing His-MBP-Plk4 expression plasmid. This work was supported by the Division of Intramural Research at the NIH/NHLBI (1ZIAHL006104) to N.M.R. G.C.R. is grateful for support from NCI P30 CA23074, NIH/NIGMS R01GFM110166, the National Science Foundation MCB1158151, and the Phoenix Friends.

Authors declare no competing financial interests.

## Supplemental materials

### Supplemental Figure Legends

**Figure S1.**
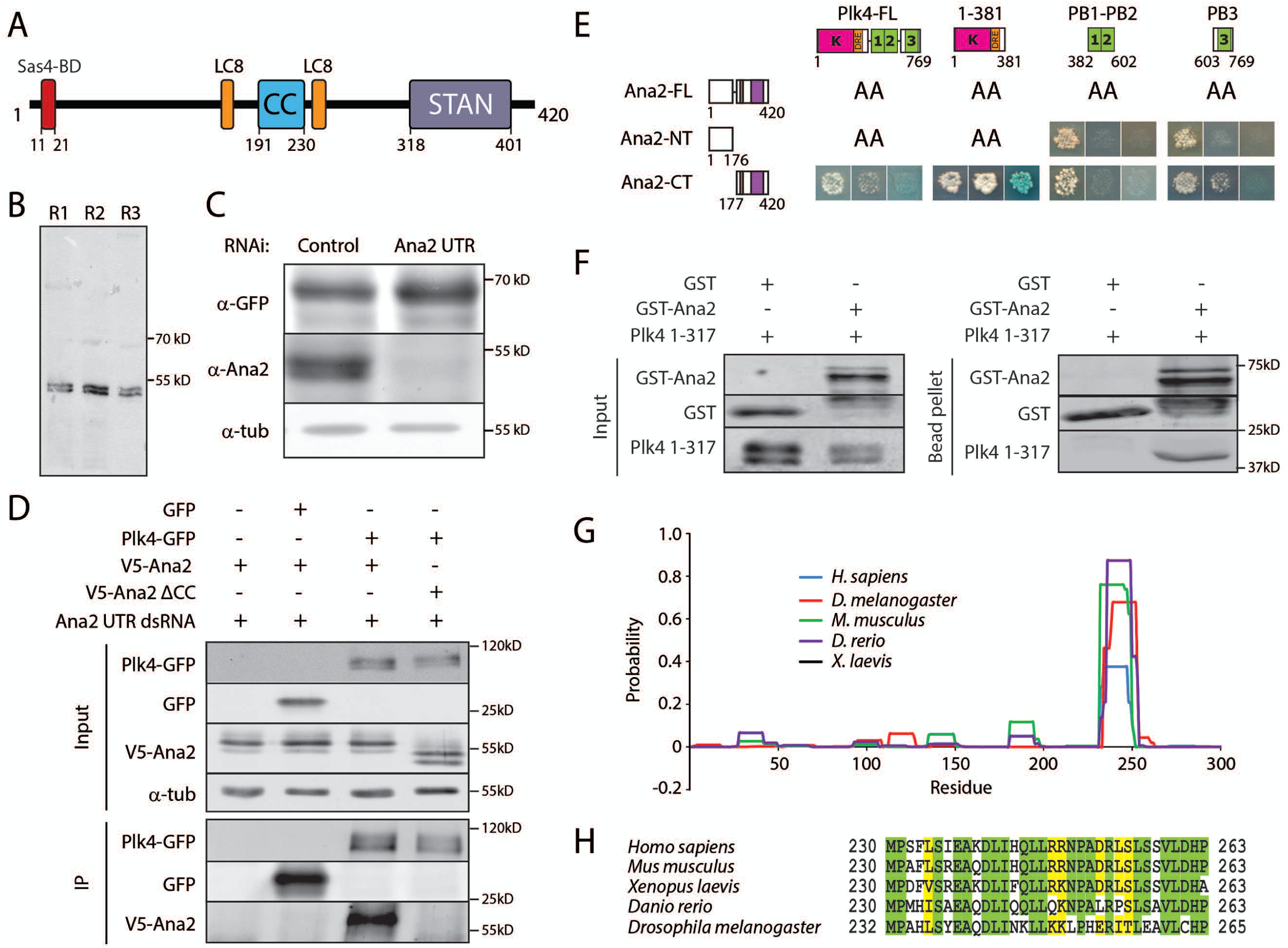
The coiled-coil domain of Ana2 binds the putative coiled-coil and Polo Box 3 (PB3) domains of Plk4. (A) Linear map of *Drosophila* Ana2 showing the functional and structural domains, including the Sas4-binding domain (red) (Cottee et al., 2013), cytoplasmic dynein light chain LC8-binding regions (amino acids 159–168 and 237–246; orange) (Slevin et al., 2014), coiled-coil region (CC, blue), and the STil/ANa2 (STAN) motif (purple). (B) The anti-Ana2 antibodies used in this study are specific. Three different rabbit polyclonal anti-Ana2 antibodies were raised against purified recombinant full-length GST-Ana2 and then affinity-purified with MBP-Ana2. The specificity of these antibodies (R1-3) on Western blots of S2 cell lysates is shown. All antibodies react with the identical ~50 kDa protein which migrates on SDS-PAGE as a tight doublet. (C) Ana2 RNAi efficiently diminishes Ana2 levels in cells. Ana2 dsRNA was generated against sequence of the 5’ and 3’ UTRs. S2 cells were treated with control dsRNA or Ana2 UTR dsRNA for 7 days. On day 5, cells were transfected with GFP-Ana2, allowed to recover for 24 hours, and then induced to express transgenic GFP-Ana2 for an additional 24 hours. Anti-Ana2 immunoblots demonstrate effective endogenous Ana2 depletion (~90%). Anti-GFP immunoblots show that Ana2 UTR RNAi does not diminish expression of exogenous GFP-Ana2. (D) Ana2 but not Ana2 ΔCC co-immunoprecipitates (IPs) with Plk4. S2 cells were RNAi treated for 7 days to deplete endogenous Ana2. On day 5, cells were co-transfected with Plk4-GFP (or control GFP) and either V5-Ana2 wild-type or V5-Ana2 ΔCC; the following day, transgene expression was induced for 24 hrs. Anti-GFP IPs were then prepared from lysates, and blots of the inputs and IPs probed for GFP, V5, and α-tubulin. (E) Ana2 interacts specifically with Plk4 1-381 and PB3 by yeast two-hybrid (Y2H) analysis. Full-length (FL) and fragments of Ana2 and Plk4 were screened by Y2H. In each image, colonies from replica plating are shown and growth indicates the presence of both bait and prey. Left panel, no selection; middle panel, growth selection on QDO; right panel, growth and color selection on DDOXA (blue color indicates an interaction). AA indicates that one or both protein fragments autoactivated the Y2H reporters on their own and could not be tested. Note that in this subfigure only, the Ana2-CT is defined as spanning residues 177-420 and contains the CC. Ana2-CT interacts strongly with Plk4 1-381 (containing the CC-DRE region) and moderately with PB3, but not with PB1-PB2. (F) In vitro GST-pulldown assays demonstrate direct binding between purified GST-Ana2 and FLAG-tagged Plk4 1-317-His_6_. Western blots of input and washed glutathione bead pellet samples were probed with anti-GST and FLAG antibodies. (G) Coiled–coil (CC) regions of Plk4 family members predicted by software from Pole Bioiformatique Lyonnaise (PBIL). The probability of CC structure was determined for the first 300 amino acids, using a 14 amino acid window and a 2.5 weight on positions ‘a’ and ‘d’. (H) Alignment of the 34 amino acid within the predicted CC encoded by Plk4 family members. Identical residues in the alignment are highlighted green, and aligned residues with similar side chains are highlighted yellow.

**Figure S2.**
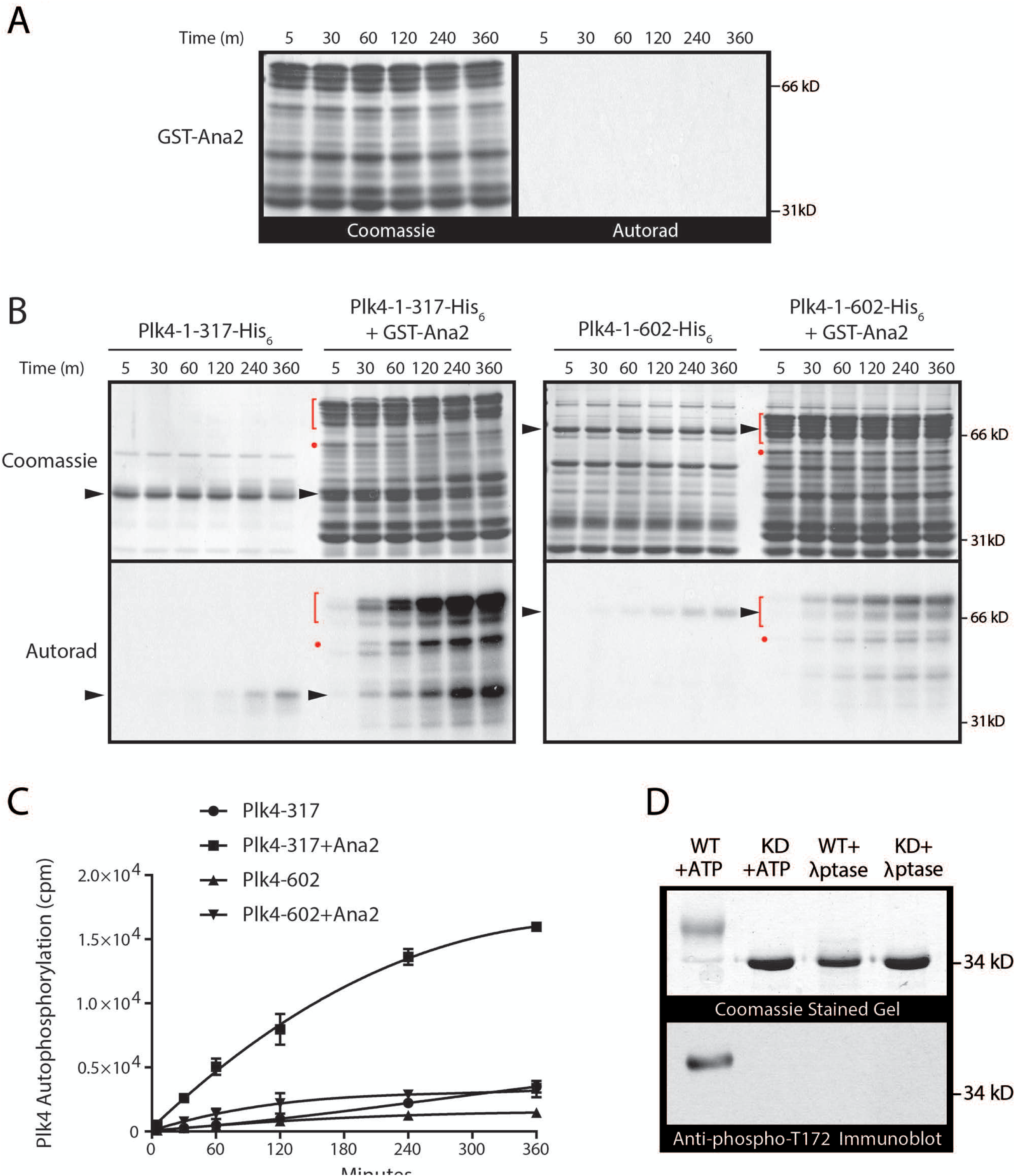
Ana2 stimulates full-length Plk4 kinase activity but not an autoinhibited Plk4 mutant lacking PB3. (A) GST-Ana2 lacks detectable kinase activity. GST-Ana2 was incubated with γ^32^P-ATP and sampled at intervals. Left, Coomassie-stained SDS-PAGE gel of samples; right, corresponding autoradiogram (exposed with the same conditions used for the autoradiograms in B). (B, C) Ana2 increases the autophosphorylation of Plk4 1-317 but not Plk4 1-602. Purified Plk4 1-317-His_6_ (317) and Plk4 1-602-His_6_ were incubated with γ^32^P-ATP and each reaction was sampled at intervals (0–360 min). Equimolar amounts of kinase were used in the reactions and loaded in each lane. (B) Top panels, Coomassie-stained gels; bottom panels, autoradiograms (exposed under identical conditions). Arrowheads mark positions of Plk4 317 and 602. (C) Graph of the scintillation counts of Plk4 bands cut from gels. n = 3; error bars show SEM. (D) Anti-phospho-Plk4 polyclonal antibody was generated to specifically recognize phospho-Thr172 (T172), an autophosphorylated residue in the Plk4 activation loop and a readout for activated kinase. Bacterially-expressed, purified Plk4 1-317-His_6_ or kinase dead (KD) Plk4 1-317-His_6_ was incubated with ATP and subsequently, either treated with λ-phosphatase or mock treated as a control. Anti-pT172 antibody specifically recognizes Plk4, but not Plk4 KD or Plk4 treated with λ-phosphatase. Coomassie-stained protein gel (top) and the corresponding immunoblot (bottom) are shown. Note that whereas KD and λ-phosphatase-treated Plk4 migrate on SDS-PAGE as a single fast-migrating non-phosphorylated species, autophosphorylated Plk4 (WT Plk4 incubated with ATP) migrates as a series of phosphorylated polypeptides, appearing as a diffuse band or even a short ladder, corresponding to Plk4 phosphorylated to different extents (Klebba et al., 2015a).

**Figure S3.**
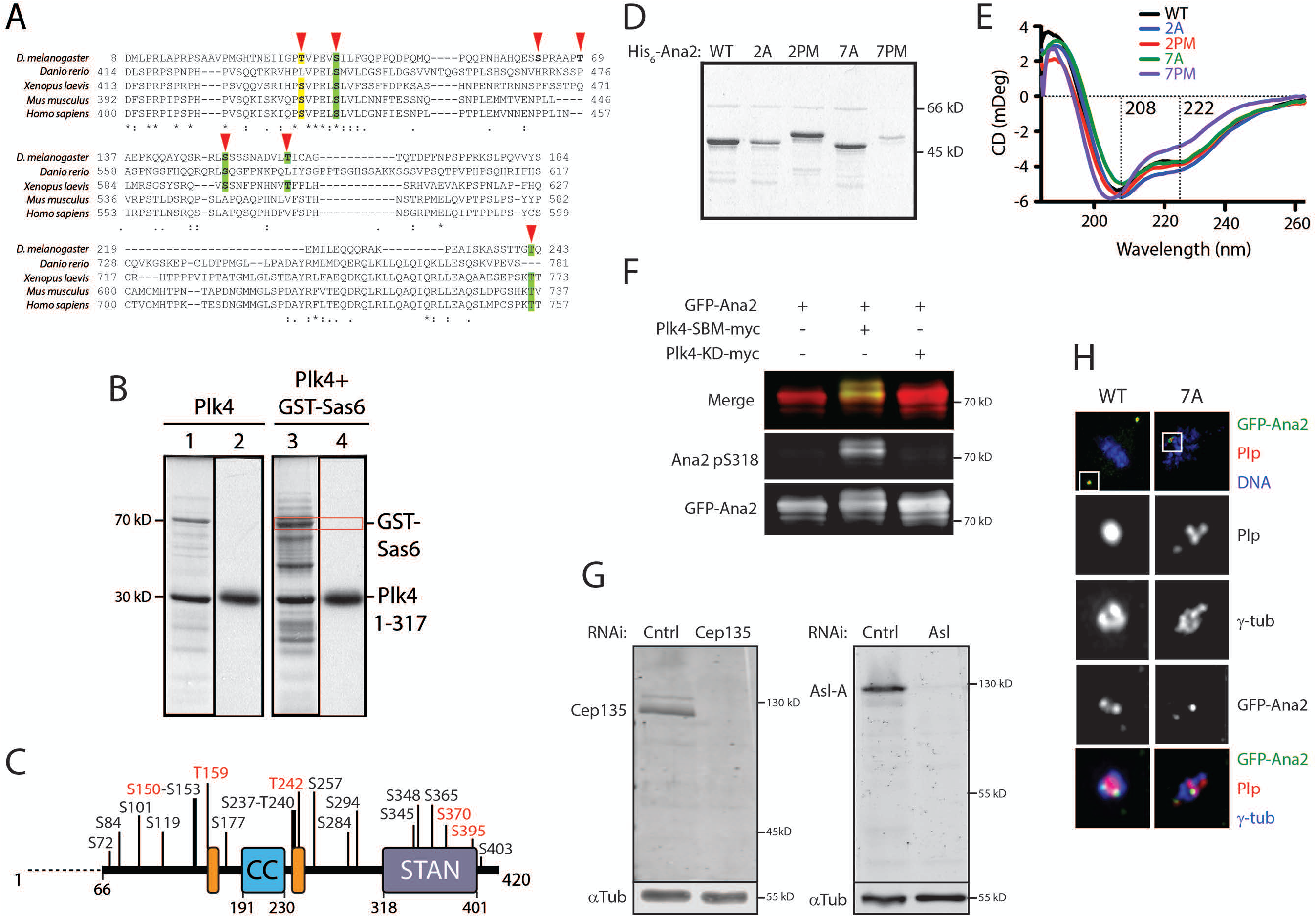
Plk4 phosphorylation of Ana2 N-terminal residues regulates centriole duplication. (A) Primary sequence alignment of Ana2/STIL family members. Identical residues are denoted with an asterisk and partially conserved residues with a colon or period below the alignment. Positions of the seven Plk4 phosphorylated residues that lie upstream of the STAN domain in fly Ana2 are indicated (red arrowheads). Five of these residues T33/S38/S150/T159/T242 show some degree of conservation across phyla. Of these five, identical residues are highlighted green, and residues with similar side chains are highlighted yellow. Lineup was obtained using Muscle Multiple alignment. (B) Plk4 does not phosphorylate Sas6 in vitro. Purified Plk4 1-317-His_6_ was incubated with γ^32^P-ATP and with purified, full-length GST-Sas6 (lanes 3, 4). Lanes 1 and 3, Coomassie-stained gel; lanes 2 and 4, corresponding autoradiograms. Plk4 autophosphorylates but does not phosphorylate GST-Sas6 (red box). (C) Linear map of Ana2 depicting in vivo phosphorylated sites identified in GFP-Ana2 immunoprecipitated from S2 cell lysates. Many of these phosphorylated residues are not targeted by Plk4 in vitro and could be targets of one or more unidentified kinases. For example, STIL is also a Cdk1 substrate (Zitouni et al., 2016). Some sites could not be precisely located: one residue within S150-S153 is phosphorylated (probably S150), and only one residue within S237-T240 is phosphorylated. Residues 1-65 were not recovered. (D) Purity of Ana2 constructs used for Circular Dichroism. GST-Ana2 WT and mutants were bacterially expressed and purified on glutathione resin. The GST tag was removed by proteolytic cleavage, and the purified protein was visualized by Coomassie staining of the SDS-PAGE gel. (E) CD spectra of full-length His_6_-Ana2 wild-type, 2A, 2PM, 7A, and 7PM constructs. Constructs show minima at 208 and 222 nm, indicative of α-helical secondary structure, consistent with secondary structure predictions, which assign α-helical structure to the tetramerization domain [verified in structure 5AL6 by Cottee et al. (2015)] and the C-terminal STAN motif, two domains that do not overlap with any of the seven phosphorylation sites investigated in this study. All mutants yielded spectra similar to wild-type Ana2, though the 7PM mutant showed a slight shift from the 208 minimum that may be due to its lower expression level which likely yielded higher levels of impurities. Representative traces are shown from data collected on two independent experimental days. (F) Immunoblots of anti-GFP-Ana2 immunoprecipitates demonstrate specificity of the pS318 antibody. Cells were co-transfected with GFP-Ana2 and inducible kinase-dead (KD) or non-degradable Slimb-binding mutant (SBM) Plk4-myc. 24 hours after transfection, cells were induced to express for 24 hours. Anti-GFP IPs were then prepared from lysates, and Western blots of the IPs probed for GFP and pS318. A merge of the GFP-Ana2 and pS318 blots is shown (upper panel). (G) The anti-Cep135 and anti-Asl N-terminal (NT) antibodies used in this study are specific. Rat polyclonal anti-Cep135 and Asl-NT antibodies were raised against purified recombinant full-length GST-tagged proteins and then affinity-purified with MBP-tagged proteins. The specificity of these antibodies on immunoblots of S2 cell lysates is shown next to Cep135 and Asl-depleted cell lysates. α-Tubulin (α-Tub) was used as a loading control. (H) Centrioles containing GFP-Ana2-7A recruit pericentriolar material during mitosis. S2 cells were transfected with the indicated GFP-Ana2 (green) construct and induced to express for 5 days. Cells were immunostained for PLP (red), γ-tubulin (green) and DNA (blue). Deconvolution micrographs show GFP-Ana2-WT in a metaphase cell and Ana2-7A in a prometaphase cell. Dashed boxes in the first row of images are shown at higher magnification in rows 2–5. Scale, 5 μm.

**Figure S4.**
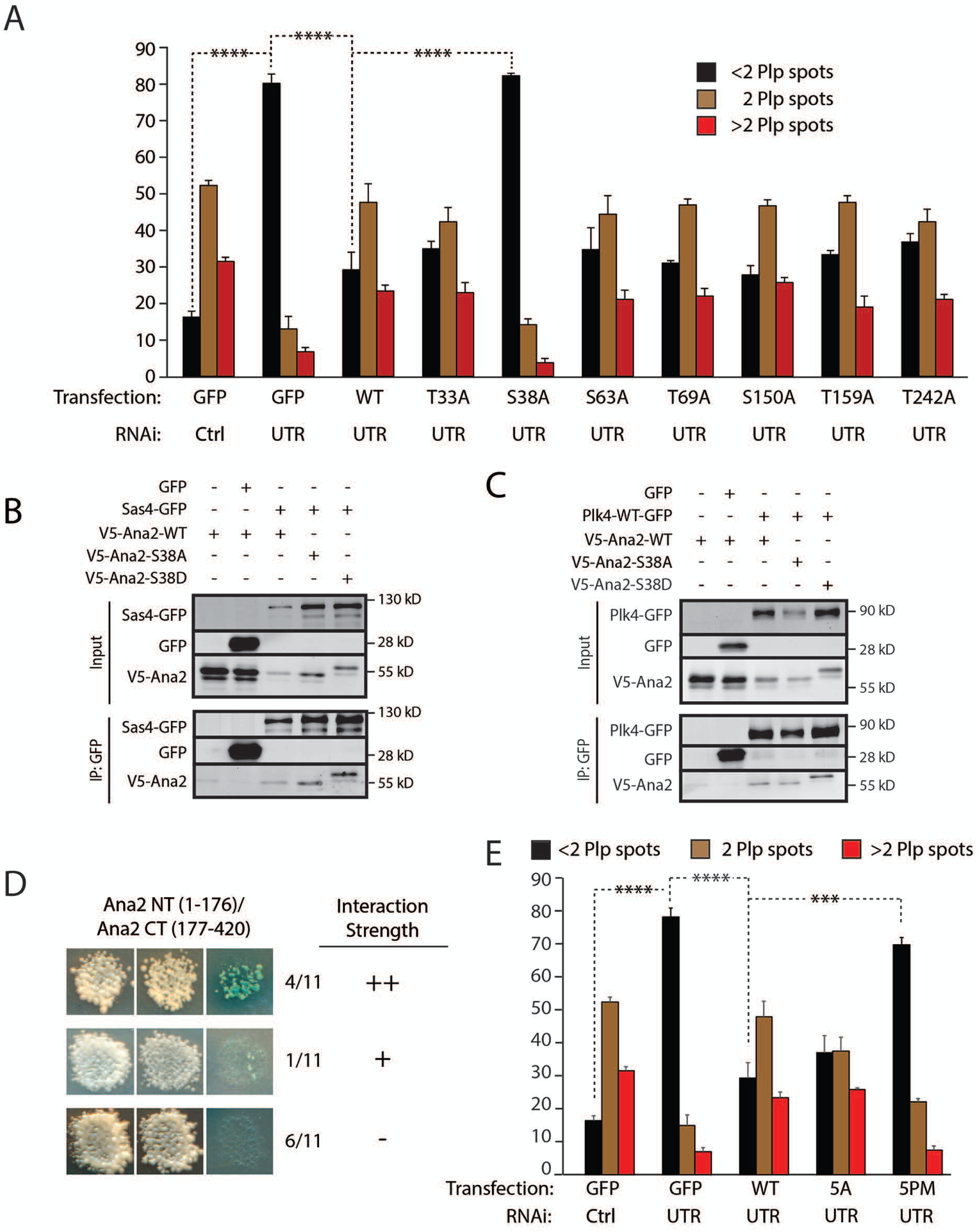
S38 phosphorylation is critical for centriole assembly, but mutation of S38 alone does not influence Sas4 or Plk4 binding. (A) A single non-phosphorylatable alanine substitution of S38 in Ana2 disrupts centriole duplication. S2 cells were control-dsRNA treated (Ctrl) or depleted of endogenous Ana2 (UTR) by RNAi for 12 days. Cells were transfected with Ana2-UTR dsRNA on days 0, 4 and 8. On days 4 and 8, cells were additionally transfected with V5-Ana2 or GFP and induced with 0.1 mM CuSO4. Cells were immunostained for PLP and Asterless to mark centrioles, and the number of centrioles per cell was counted. n = 100 cells in each of three experiments. Asterisks indicate significant differences. Error bars, SEM. (B, C) Phospho-mutants of S38 in Ana2 do not affect binding to Sas4 (B) or Plk4 (C). S2 cells were depleted of endogenous Ana2 by RNAi for 7 days. On day 5, cells were co-transfected with the indicated inducible V5-Ana2 construct and either Sas4-GFP (B) or Plk4-GFP (C), and the next day induced to express for 24 hours. Anti-GFP IPs were then prepared from lysates, and Western blots of the inputs and IPs probed for GFP and V5. (D)Ana2 NT interacts specifically with Ana2 CT by yeast two-hybrid (Y2H) analysis. Fragments of Ana2, NT (1-176) and CT (177-420) were screened by Y2H. In each image, colonies from replica plating are shown and growth indicates the presence of both bait and prey. Left panel, no selection; middle panel, growth selection on QDO; right panel, growth and color selection on DDOXA (blue color indicates an interaction). Note that in this subfigure only, the Ana2-CT is defined as spanning residues 177-420 and contains the CC. Ana2-CT interacts strongly with Ana2-NT in 4 out of 11 interactions, weakly in 1 out of 11, and no interaction was detected in 6 out of 11 interactions. (E) Ana2-5PM does not rescue centriole duplication. S2 cells were control-dsRNA treated (Ctrl) or depleted of endogenous Ana2 (UTR) by RNAi for 12 days. Cells were transfected with Ana2-UTR dsRNA on days 0, 4 and 8. On days 4 and 8, cells were additionally transfected with V5-Ana2 or GFP and induced with 0.1 mM CuSO4. Cells were immunostained for PLP and Asterless to mark centrioles, and the number of centrioles per cell was counted. n = 100 cells in each of three experiments. Asterisks indicate significant differences. Error bars, SEM.

### Supplemental Table

**Table S1.**
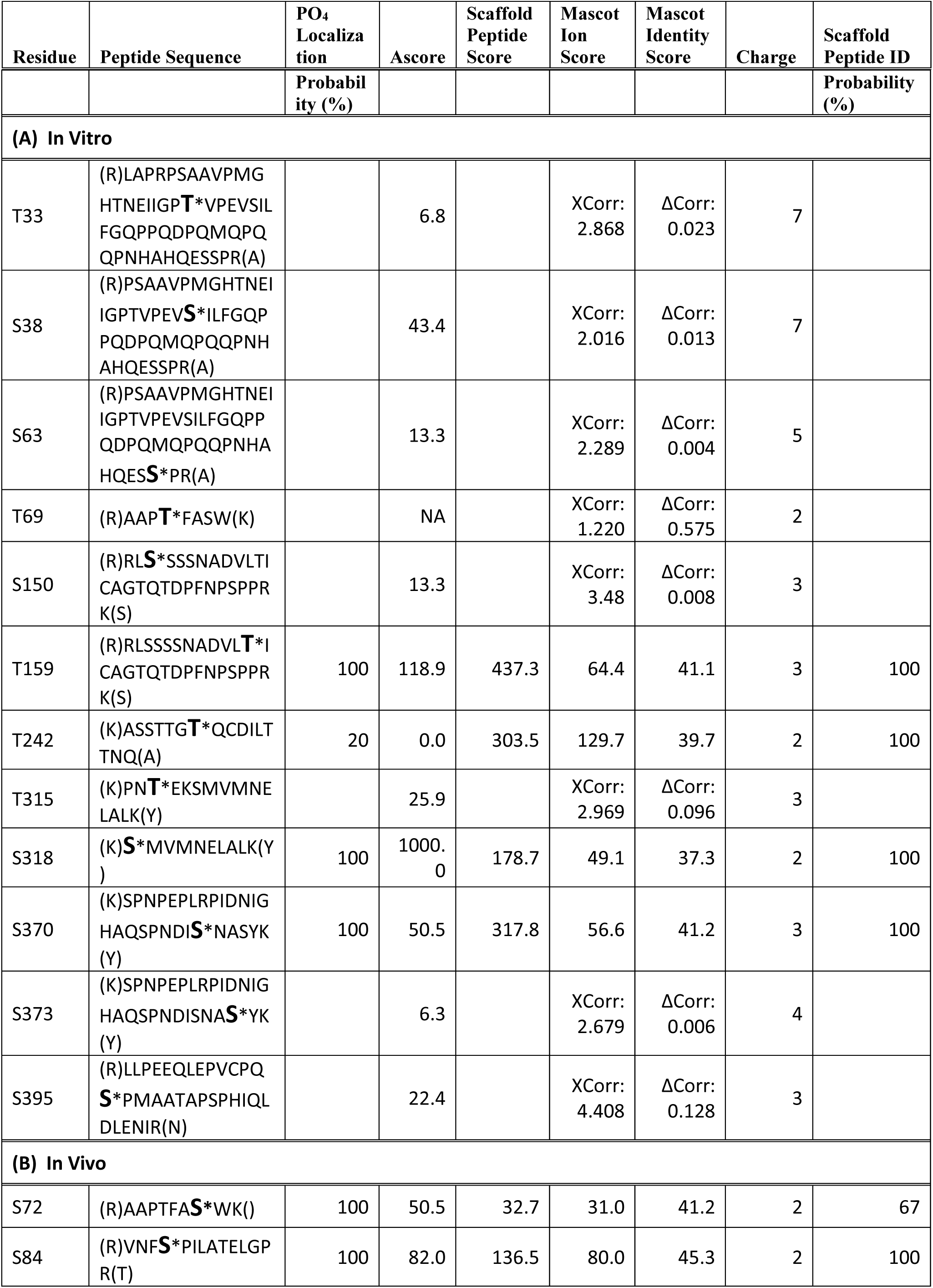
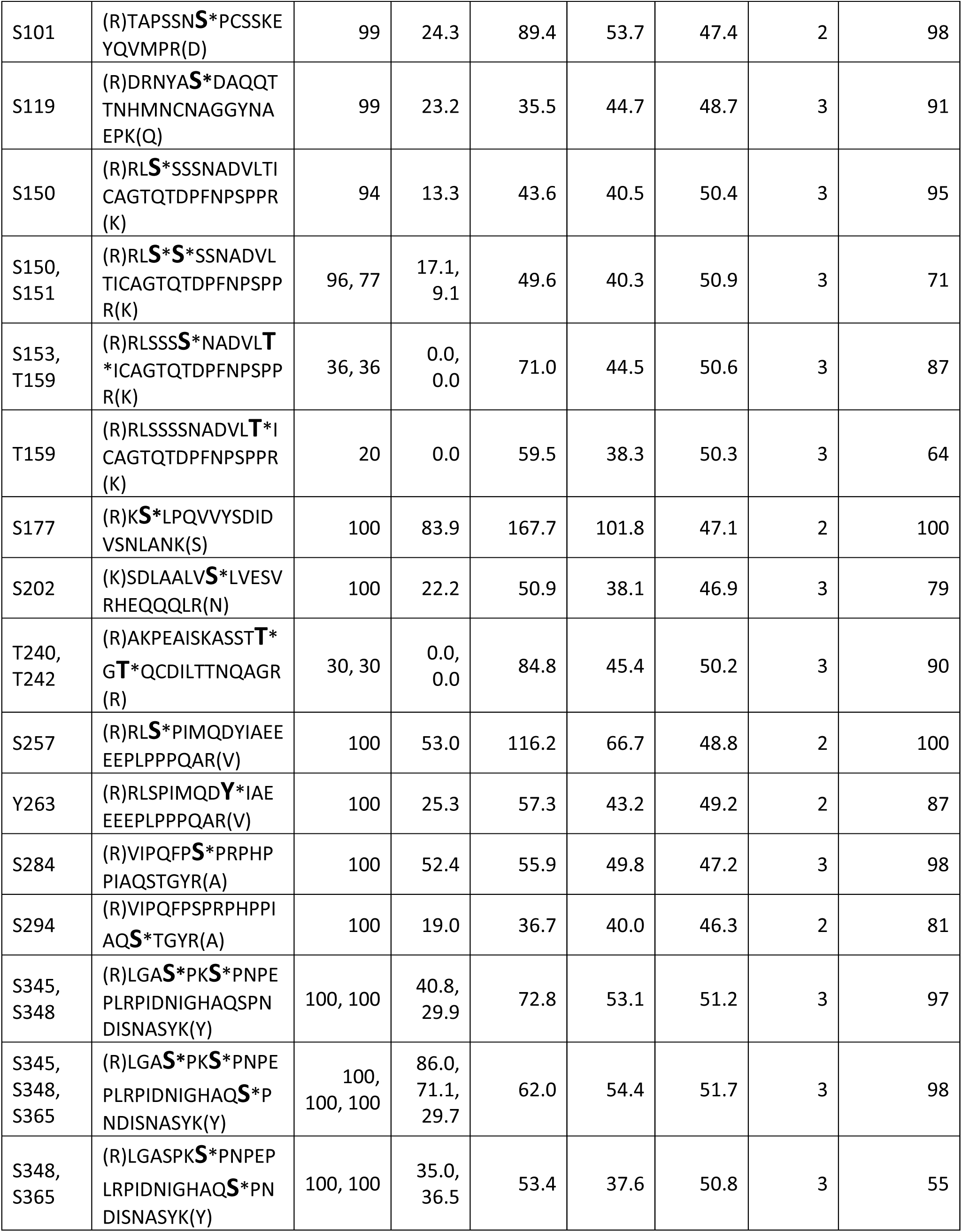
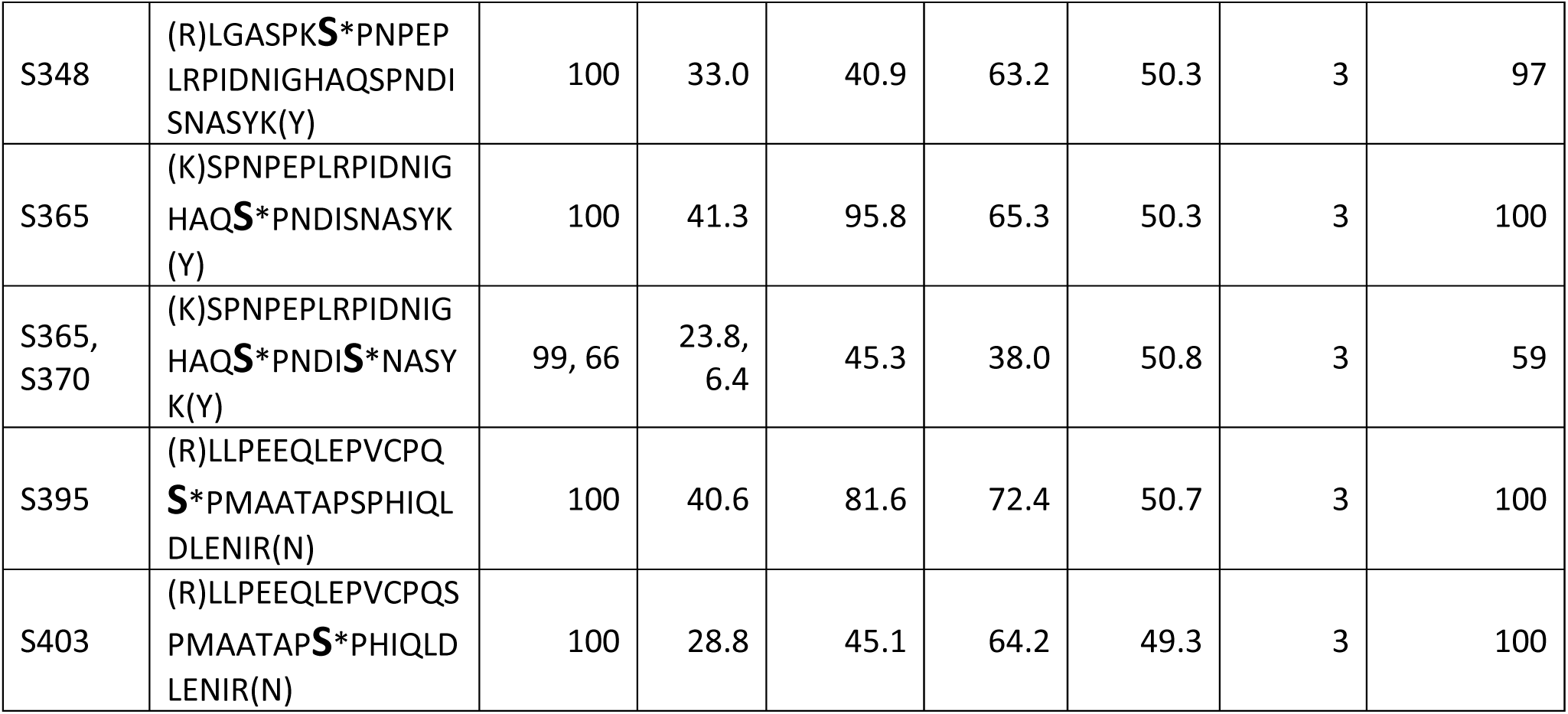
In vitro and in vivo phosphorylated residues of Ana2. Phosphorylated residues of Ana2 are indicated with bold, large font followed by an asterisk. Ana2 samples were analyzed by LC-MS/MS by three different proteomics facilities, and either Mascot or Sequest program was used to identify peptides from a *D. melanogaster* protein database (NCBI). Xcorr and ΔXcorr values were obtained from Sequest (and, when available, Ascores were supplied by Taplin Facility, Harvard). Otherwise, displayed values were obtained from ScaffoldPTM (Proteome Software). (A) In vitro phosphorylated residues of Ana2 were identified by first incubating purified, bacterially-expressed *D. melanogaster* Plk4 kinase domain (amino acids 1-317) and GST-Ana2 with MgATP as described in Methods. Ana2 was then resolved by SDS-PAGE, and Coomassie-stained bands were processed (e.g., proteolyzed) for LC-MS/MS. (B) Phosphorylated residues of transgenic Ana2 purified from lysates of transfected S2 cells were similarly identified using LC-MS/MS analysis of processed, SDS-PAGE resolved Ana2.

